# Neural tube development depends on notochord-derived Sonic hedgehog released into the sclerotome

**DOI:** 10.1101/639831

**Authors:** Nitza Kahane, Chaya Kalcheim

## Abstract

Sonic hedgehog (Shh), produced in notochord and floor plate, is necessary both for neural and mesodermal development. To reach the myotome, Shh has to traverse the sclerotome. By loss and gain of Shh function, and floor plate deletions, we report that sclerotomal Shh is also necessary for neural tube development. Reducing the amount of Shh in sclerotome by membrane-tethered hedgehog-interacting protein or by Patched1, but not by dominant active Patched, decreased motoneuron numbers while also compromising myotome differentiation. These effects were a specific and direct consequence of reducing Shh. In addition, grafting notochords in a basal, but not apical location vis-a-vis the tube, profoundly affected motoneuron development, suggesting that initial ligand presentation occurs at the basal side of epithelia corresponding to the sclerotome-neural tube interface.

Collectively, our results reveal that the sclerotome is a potential site of a Shh gradient that coordinates development of mesodermal and neural progenitors.

**Summary statement:** Shh that transits through the sclerotome is presented to the neuroepithelium from its basal aspect to affect motoneuron development.

## Introduction

The morphogen Sonic hedgehog (Shh) plays fundamental roles in the development of both neural tube (NT) and somites (Borycki et al., 1998; Briscoe, 2009; Cairns et al., 2008; Ericson et al., 1997a; Ericson et al., 1997b; Gustafsson et al., 2002). Its signaling is initiated by binding of the proteolytically processed and lipid modified ligand to the transmembrane receptor Patched (Ptc), that represses the pathway in the absence of ligand (Goodrich et al., 1997; Hidalgo and Ingham, 1990; Ingham et al., 1991; Johnson et al., 1996). Ligand binding to Ptc abrogates its repressive effect on Smoothened (Smo), a key effector essential for canonical Hedgehog signal transduction (van den Heuvel and Ingham, 1996). The repressive role of Ptc correlates with its localization in the primary apically located cilium, that functions as a signal transduction compartment (Caspary et al., 2007; Rohatgi et al., 2007). Binding of Shh to Ptc removes the latter from the cilium, thereby allowing Smo to enter and propagate the signal further downstream (Milenkovic et al., 2009; Rohatgi et al., 2009) to regulate Gli transcription factor activity (Briscoe and Therond, 2013; Ribes and Briscoe, 2009).

Shh signaling is highly regulated by negative and positive modulators. Ptc1, Hedgehog interacting protein (Hhip) and Gli1 are direct targets of Shh and the former two also inhibit its activity (Chuang and McMahon, 1999; Ingham and McMahon, 2001). Sulfatase1 (Sulf1) is a known Shh target (Dhoot et al., 2001) and the co-receptors Boc, Gas and Cdo (Allen et al., 2011; Izzi et al., 2011) all enhance ligand activities and are expressed in the NT and/or developing mesoderm (Kahane et al., 2013).

It is well established that at early stages following neurulation, Shh secreted by the notochord (No) induces distinct ventral cell identities in the overlying NT by a mechanism that depends on relative concentrations and duration of exposure (Briscoe and Small, 2015; Dessaud et al., 2010; Stamataki et al., 2005). Moreover, its activity continues beyond this stage to further regulate cell proliferation, survival and differentiation (Cayuso et al., 2006; Charrier et al., 2001). No-derived Shh is also involved in mesoderm patterning (Borycki et al., 1998; Cairns et al., 2008). A ventro-dorsal activity gradient of Shh/Gli signaling in sclerotome was directly visualized using an *in vivo* reporter in mice (Kahane et al., 2013). In addition, in chick embryos, Shh spreads from the midline through the sclerotome to reach the dermomyotome (DM). There it promotes terminal myogenic differentiation of both epaxial and hypaxial DM-derived progenitors and maintains epitheliality of DM cells (Kahane et al., 2013). Notably, in both floor plate (FP) and myotome, the activities of Shh are transient and cells become refractory to the ligand, a mechanism that allows dynamic phase transitions to take place within these systems (Cruz et al., 2010; Kahane et al., 2013).

Because Shh is important for the development of both NT and mesoderm, two functionally interconnected systems, the question arises whether the effects of Shh on either tissue are independent of each other or interrelated. Furthermore, does the NT receive Shh directly from the producing sources (No and FP), or given the ligand is released into the mesoderm, can the latter serve as an “en passant” pathway from which Shh affects aspects of both NT and mesoderm development?. Answering these questions is of utmost significance both for better understanding the mechanism of Shh activity and for achieving an integrated molecular view of regional development.

In this study we report that, in addition to affecting muscle development, reducing the amount of Shh in the sclerotome by Hhip1 also significantly reduces motoneuron numbers in NT. Notably, similar effects are monitored when a membrane-tethered version of Hhip1 (Hhip:CD4), unable to be released from cells, is locally electroporated into the sclerotome. The observed phenotypes are a specific and direct consequence of Shh depletion as they are rescued by excess Shh, direct Shh targets such as *Hhip1* and *Gli1* mRNAs are reduced, and its effects are not mediated through other signaling pathways. Most importantly, the effects of Hhip:CD4 are phenocopied by electroporation of the transmembrane receptor Ptch1 but not by constitutively active PTC^Δloop2^ which does not recognize the ligand. In addition, by gain and loss of Shh function, and by FP deletions, we show that the sclerotome, but not the NT itself, constitutes a dynamic substrate of No-derived Shh that acts both on motoneuron as well as on myotome development. Furthermore, grafting No fragments adjacent to the basal, sclerotomal side of the NT profoundly affects its development when compared to apical grafts. A similar basal grafting with respect to the DM significantly enhances myotome formation, suggesting a general need for initial ligand presentation at the basal side of epithelia. Together, our results uncover the sclerotome as a novel pathway through which No-derived Shh disperses to promote aspects of neural development.

## Results

### Reduction of Shh in sclerotome by Hhip1 affects both myotome and motoneuron differentiation

To investigate possible interactions between neural and mesodermal progenitors mediated by Shh, electroporations were performed in avian embryos aged 23-25 somite pairs at the level of epithelial somites prior to myotome and sclerotome formation. This is the earliest time point at which the prospective sclerotome can be faithfully attained by focal electroporation. In this region, the NT is composed of proliferative cells (Kahane and Kalcheim, 1998) and neural patterning is already apparent and ongoing, as evidenced by expression of the positive Shh targets *Nkx2.2, Olig2, Nkx6.2 and Nkx6.1* (Supplem. Fig. S1, A-D). However, differentiation into Hb9-expressing motoneurons has not yet occurred at this stage (Supplem. Fig. S1, E) and only starts about 10 hr later at the level of somites 11-12 located rostral to the last segmented pair of somites (Supplem. Fig. S1, F, arrows). Hence, the timing of manipulations of Shh activity corresponds to the transition of specified proliferative progenitors into differentiated motoneurons (Ericson et al., 1996). Hence, we focused on effects of altering Shh activity in mesoderm on the survival, proliferation and differentiation of motoneurons, all Shh-dependent events occurring at a post-patterning phase.

In a previous study, we reported that Shh traversing the sclerotome is necessary for myotome differentiation, as missexpression of the high affinity and selective Shh antagonist Hhip1 in the sclerotome resulted in smaller myotomes expressing desmin accompanied by a corresponding accumulation of Pax7-positive progenitors [(Kahane et al., 2013) and Fig. 1, A,B]. Here, we report that the hemi-NT facing the transfected mesoderm was also affected, exhibiting a 40% reduction in the number of Hb9+ motoneurons when compared to controls that received GFP alone (asterisks in Fig. 1A’,B’, C, p<0.001, N= 12 and in both control and treated embryos). This is a significant effect given that electroporation is a mosaic technique that attains only a fraction of cells in the transfected domain thus causing a reduction rather than a total depletion of the ligand. Moreover, a ventral expansion of the Pax7-positive domain was frequently observed (arrows in Fig. 1A,B), likely due to a reduced size of the ventral extent of the transfected hemi-NT (see Fig.2).

**Fig. 1.**
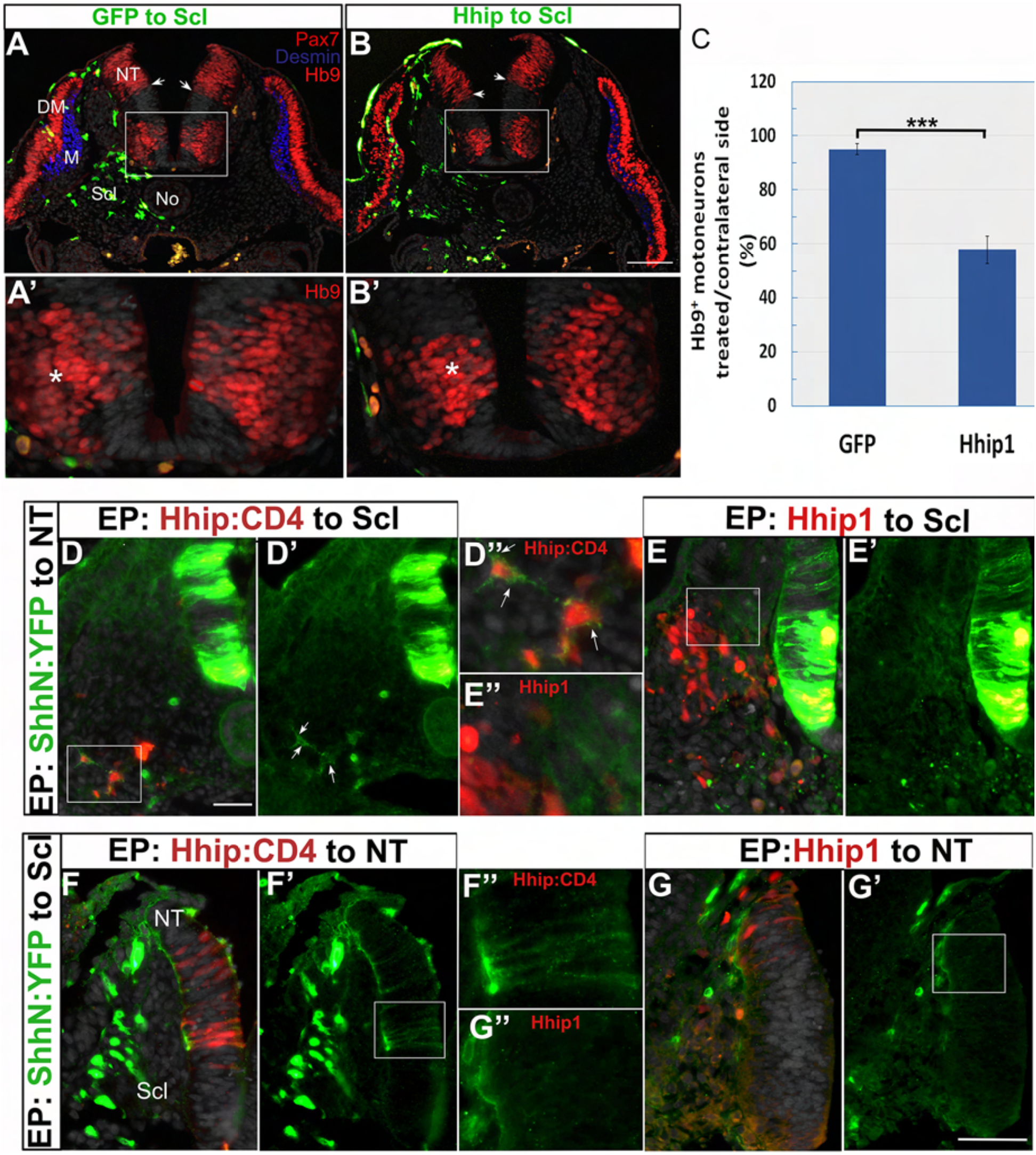
Reduction of Shh in sclerotome by Hhip1 affects both myotome and motoneuron differentiation. (A, B) Electroporation of control GFP (A) or Hhip1/GFP (B) to the prospective sclerotome. A day later a reduction in myotome size (blue desmin staining) is apparent adjacent to the transfected cells in B when compared to the control side and to control GFP (A). In addition, note the ventral expansion of the Pax7+ domain in B (small arrows) when compared to the control side and to control GFP. (A’, B’) Higher magnification of the insets in A,B, respectively depicting a reduction of Hb9+ motoneurons in B upon Hhip1 treatment when compared to the control side and to control GFP (A’). Asterisks (*) denote motoneurons adjacent to the electroporated sclerotomes. (C) Quantification of Hb9+ motoneurons (***, p<0.001). (D-D”) Double electroporation of ShhN:YFP to NT and Hhip1:CD4 to sclerotome. Note that sclerotomal cells missexpressing Hhip:CD4 (red) are decorated by Shh (green) immunolabeling (D’, D” arrows). (E-E”) Double electroporation of ShhN:YFP to NT and Hhip1 to sclerotome. No labeling of Shh (green) is apparent in sclerotomal cells missexpressing Hhip1 (red). D” and E” are high magnifications of insets in D and E. (F-F”) Double electroporation of ShhN:YFP to sclerotome and Hhip1:CD4 to NT. Note labeling of Shh (green) in both the basement membrane as well as along neuroepithelial cells missexpressing Hhip:CD4 (F’, F”). F” is a higher magnification of the inset in F/F’ where only ShhN:YFP (green) is shown to highlight that Shh co-localizes with the red, Hhip1:CD4-missexpressing cells, in the neuroepithelium (F). (G-G”) Double electroporation of ShhN:YFP to sclerotome and Hhip1 to NT. Note labeling of Shh (green) in basement membrane but no co-staining of neuroepithelial cells missexpressing Hhip1. G” is a high magnifications of the inset in G’. Abbreviations, DM, dermomyotome; M, myotome; NT, neural tube; No, notochord; Scl, sclerotome. Bars=50 μM.

**Fig.2.**
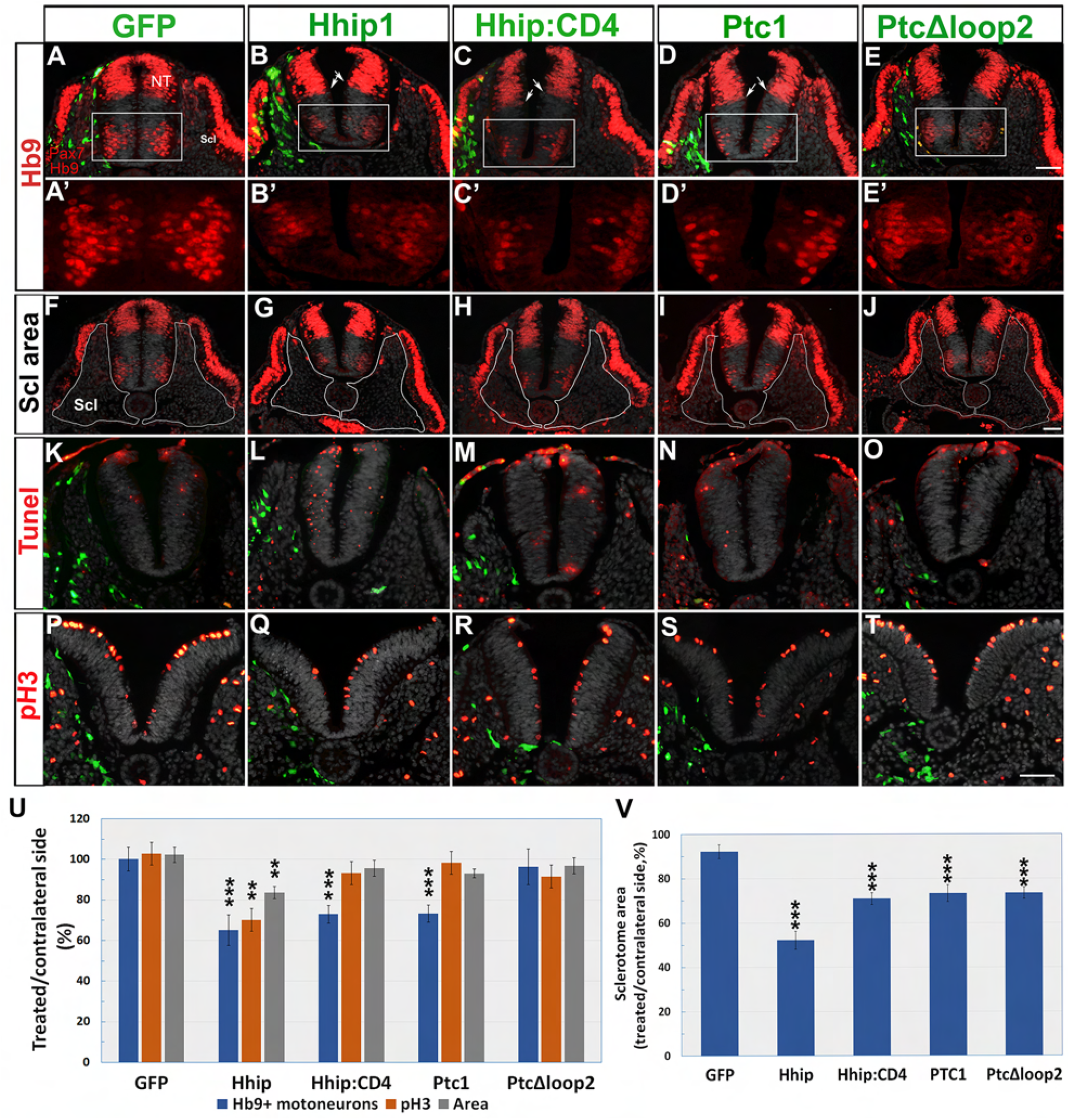
Electroporation of Hhip:CD4 and Ptc1 into sclerotome inhibit motoneuron differentiation without significantly affecting progenitor proliferation or survival. (A-E’) Electroporation of depicted plasmids (green) to the sclerotome followed a day later by immunostaining for Pax7 and Hb9. A’-E’ represent higher magnifications of the boxed regions in A-E. Note that all plasmids except PTC^Δloop2^ reduced the number of motoneurons adjacent to the transfected sclerotomes. Arrows in B-D show a slight ventralization of the ventral domain of Pax7. (F-J) The same sections as in A-E in which the sclerotomes were delineated by a white lines. (K-O) Tunel staining (red). Note weak immunostaining of electroporated plasmids (green) because no anti-GFP antibodies were implemented to enable better visualization of apoptotic nuclei. Only Hhip1 exhibited numerous Tunel+ nuclei adjacent to the transfected sclerotome. Ectodermal staining reflects non-specific reactivity often seen in peripheral tissues. (P-T) staining of mitotic nuclei with anti pH3 (red). (U) Quantification of motoneurons, mitotic nuclei and area. (V) Quantification of the relative area of electroporated sclerotomes based on sections such as those shown in F-J. Abbreviations, NT, neural tube; Scl, sclerotome. *p<0.05, **p<0.03, ***p<0.01. Bar=50μM.

These results can be explained by the sclerotome constituting a substrate through which Shh disperses and from which the ligand is provided to both mesodermal and neural progenitors. This would imply that the NT does not only receive the ligand directly from No and/or FP. Alternatively, they might result from Hhip1 moving from the transfected mesoderm towards the NT. Indeed, in spite of the initial findings that Hhip1 is a transmembrane glycoprotein with cell autonomous functions (Bishop et al., 2009; Chuang and McMahon, 1999), it has recently been reported that Hhip1 is a secreted Shh antagonist able to exert long range effects on Shh signaling (Holtz et al., 2015; Kwong et al., 2014). This is compatible with the possibility that the transfected sclerotomal Hhip1 crosses into the NT or at least localizes to its surrounding basement membrane (Holtz et al., 2015).

### Differential behavior of secreted Hhip1 compared to membrane-tethered Hhip1:CD4 vis-à-vis Shh

To discriminate between the above possibilities, we produced a Hhip:CD4 plasmid encoding for a membrane-tethered version of Hhip1, unable to undergo secretion (Holtz et al., 2015; Kwong et al., 2014). First, we asked whether NT or sclerotomal cells missexpressing either Hhip1 or Hhip:CD4 are able to sequester Shh. To this end, we implemented a plasmid encoding the N-terminus of Shh fused in frame to YFP (ShhN:YFP). ShhN:YFP is able to undergo palmitoylation but not addition of a cholesterol moiety, a property that enables free movement of the mutant protein when compared to native Shh (Beug et al., 2011). When electroporated to the sclerotome, secreted ShhN:YFP was apparent along the basement membrane of the NT where it co-localized with laminin, yet no fluorescent signal was detected along the neuroepithelial cells (Supplem. Fig. S2, A-C, arrows).

When double electroporations of ShhN:YFP to the NT and Hhip:CD4 to sclerotome or vice-versa were performed, the Hhip:CD4-transfected sclerotome or NT progenitors, respectively, were decorated with ShhN:YFP, demonstrating that Hhip:CD4 binds and immobilizes the ligand on the surface of the expressing cells (Fig. 1D-D”,F-F”). In contrast, no such co-localization could be observed in either sclerotome or NT upon double electroporation of ShhN:YFP and Hhip1 (Fig. 1, E-E”, G-G”) consistent with the notion that native Hhip1 is a secreted protein.

In addition, we monitored the expression of endogenous Shh protein upon transfection of Hhip1 or Hhip:CD4 to the sclerotome. Shh immunoreactive protein was evident both intracellularly as well as associated with the cell membranes of the FP and No likely exposed to their external surface (Supplem. Fig. S2, D-F’). Electroporation of control GFP and of Hhip:CD4 had no effect on Shh immunoreactive protein in either the No or FP (Supplem. Fig. S2, D-E’). In contrast, missexpression of Hhip1 markedly reduced Shh levels in both structures unilaterally adjacent to the transfected cells; this reduction was mainly apparent at their basal sides closer to the transfected Hhip1 (Supplem. Fig. S2, arrows in F’). This effect may result from Hhip1 masking antibody binding to the ligand as the 5E1 Shh antibody and Hhip1 bind to the same pseudo-active site on the Shh molecule (Bishop et al., 2009; Maun et al., 2010). Together, the above data confirm that both Hhip as well as Hhip:CD4 bind Shh, but, in contrast to Hhip:CD4, Hhip1 is secreted to adsorb Shh far from the producing cells.

### The effects of Hhip:CD4 in the NT resemble those observed with other Shh inhibitors

Next, we employed the NT to examine the specificity of Hhip:CD4 relative to other, known inhibitors of the Shh pathway. Electroporation of Hhip:CD4, like that of Hhip1, Ptc1 or PTC^Δloop2^ to hemi-NTs, significantly reduced the number of Hb9+ motoneurons and that of pH3+ mitotic nuclei while enhancing cellular apoptosis. Furthermore, the total area of the transfected hemi-NTs, that reflects overall changes in cell number (see Methods), was significantly smaller in all treatments when compared to control GFP (Supplem. Fig. S3, N=4 for all treatments, *p<0.05, **p<0.03, ***p<0.01). The observed effects on motoneuron numbers could thus result from reduced cell differentiation or, indirectly, from effects on progenitor proliferation or survival. These data confirm that Shh acts both as a mitogen and survival factor in the NT (Cayuso et al., 2006; Charrier et al., 2001). Most importantly, they demonstrate that Hhip:CD4, by acting like Hhip1, Ptc1 or PTC^Δloop2^, is a specific tool to abrogate Shh activity.

### Local depletion of Shh activity by Hhip:CD4 or Ptc1 in sclerotome inhibits motoneuron differentiation in the NT

Next, we addressed the question whether the effects originally observed across tissues (e.g; between NT and mesoderm) with secreted Hhip1 (Fig.1), can be mimicked by missexpression of two different Shh inhibitors, Hhip:CD4 and the Shh receptor Ptc1, both membrane associated molecules.

We confirmed that electroporation of Hhip1 to sclerotome caused significant effects in the NT adjacent to the transfected mesoderm, as expected from a secreted molecule. These included a reduction in the number of Hb9+ motoneurons (Fig. 2B,B’, N=8, p<0.01), a decreased number of pH3+ mitotic nuclei (Fig.2Q, N=4, p<0.03), enhanced apoptosis (Fig.2, L, N=4) and an overall decrease in the area of the respective hemi-NT (Fig. 2U, N=4, p<0.03) when compared to controls (N=5, Fig.2 A,A’,K,P,U).

Notably, transfection of both Hhip:CD4 and Ptc1 also significantly affected motoneuron numbers (N=38, p<0.01 and N=7, p<0.01, respectively, Fig. 2C,C’D,D’,U), yet had a mild but non statistically significant effect on cell proliferation (N=4 and 7), survival or total hemi-NT area (N=4 and 6, respectively) opposite the treated sclerotomes (Fig.2 M,RN,S,U). Thus, reduction of Shh activity in mesoderm mainly affects motoneuron differentiation, contrasting with Shh abrogation in the NT where all parameters were considerably compromised (Supplem. Fig. S3). The finding that motoneuron differentiation is more sensitive to a reduced amount of ligand, indicates that progenitor proliferation, survival and motoneuron differentiation are separable processes that depend upon different Shh concentrations.

As a control for Ptc1 activity, we electroporated PTC^Δloop2^ that is unable to bind circulating Shh and acts cell autonomously to inhibit its signaling. PTC^Δloop2^, like Hhip1, Hhip:CD4 and Ptc1 adversely affected the size of the electroporated sclerotomes when compared to the intact contralateral ones (Fig.2F-J,V, N=5, p<0.01), altogether demonstrating that Shh traversing the sclerotome is necessary for proliferation and/or survival of these progenitors. In contrast, PTC^Δloop2^ had no significant effect on either proliferation, survival or total area of adjacent NT cells (Fig.2O,T,U). As expected, unlike Ptc1, PTC^Δloop2^ had no effect on motoneurons (Fig.2E,E’,U) suggesting that reduced sclerotomal size is not sufficient to account for the observed loss of motoneurons.

It is worth mentioning that Hhip1, Hhip:CD4 and Ptc1 also caused a slight ventralization of Pax7 expression (arrows in B-D) which was not apparent upon treatment with PTC^Δloop2^ (Fig.2E).

The specificity of Hhip:CD4 was further tested by examining *Olig2*, an earlier marker of motoneuron progenitors. The expression domain of *Olig2* was reduced adjacent to the electroporated sclerotome (Supplem. Fig. S4). This indicates that depletion of Shh in sclerotome also affects ongoing specification of motoneuron progenitors, a process that already begun before the time electroporations were performed (Supplem. Fig. S1).

In addition, while sclerotomal missexpression of Hhip:CD4 significantly affected the number of Hb9+ cells, co-treatment of Hhip:CD4 with Shh rescued the effect of Hhip:CD4 back to control levels, further suggesting that Hhip:CD4 abrogates Shh activity. Furthermore, Shh alone significantly enhanced motoneuron differentiation (Fig. 3, N=12, 26, 8 and 16, for control GFP, Hhip:CD4, Hhip:CD4+Shh and Shh alone, respectively, p<0.001).

**Fig. 3.**
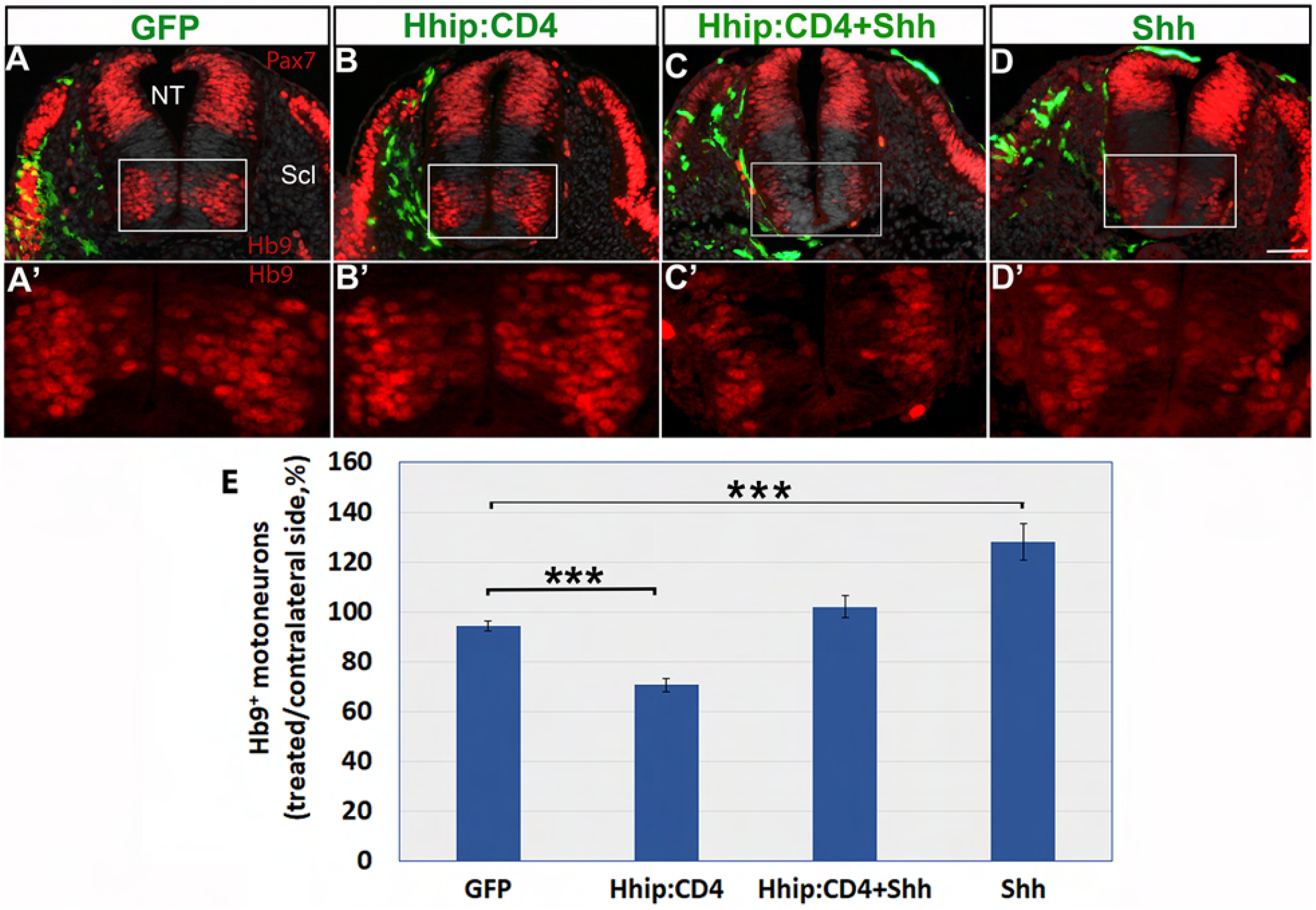
Depletion of Shh activity by Hhip:CD4 in sclerotome inhibits motoneuron differentiation and is rescued by co-treatment with Shh. (A-D) Electroporation of the depicted plasmids to sclerotome (Scl) (green). Hhip:CD4 to Scl reduces motoneuron numbers compared to control GFP, whereas co-transfection with Shh rescues the effect. A’-D’ are higher magnifications of the insets in A-D. (E) Quantification of Hb9+ motoneurons, ***p<0.001. Bar=50 μM.

### Local depletion of Shh activity by Hhip:CD4 or Ptc1 in NT inhibits myotome differentiation in mesoderm

Next, we examined whether reduction of Shh in the NT influences myotomal size. Electroporation of control GFP to hemi-NTs had no effect of myotome size (N=14) whereas Hhip1 and Hhip:CD4 caused a significant decrease in the size of the desmin+ myotomes adjacent to the transfected side of the neuroepithelium (N=3,10, respectively, p<0.001, Fig. 4, A-C, F). Electroporation of Ptc1 exhibited a similar effect (N=16, p<0.001); in contrast, PTC^Δloop2^ revealed no reduction in myotome size (N=6, Fig. 4D,E,F).

**Fig. 4.**
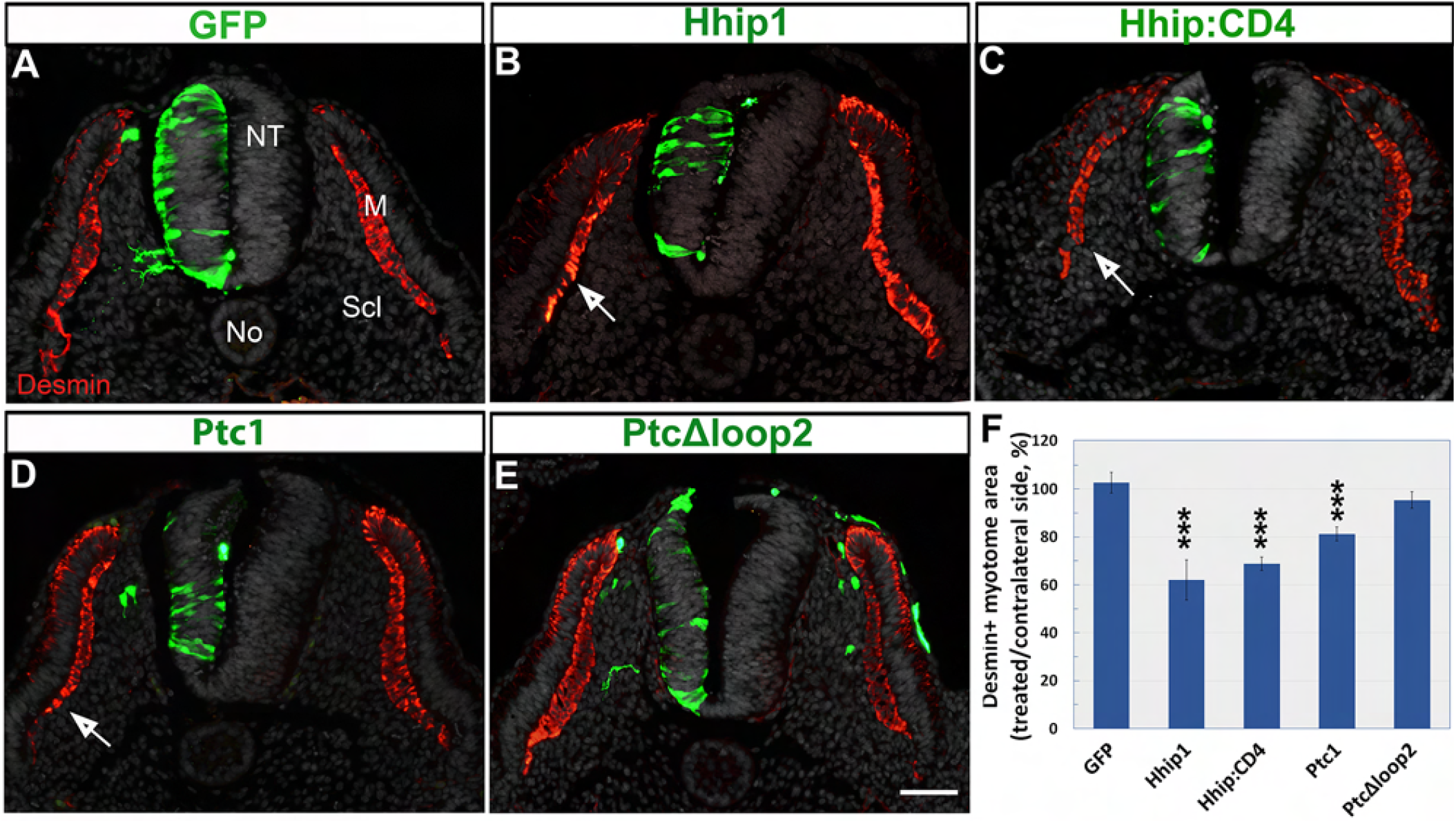
Electroporations of Hhip, Hhip:CD4 or Ptc1, but not of PTC^Δloop2^ into the neural tube, reduce myotome size. (A-E) Electroporation of the depicted plasmids (green). Hhip1, Hhip:CD4 and Ptc1 reduce the size of adjacent desmin+ myotomes (arrows, red) compared to control GFP yet PTC^Δloop2^has no significant effect. (F) Quantification of myotome size. ***p<0.001. Bar=50 μM.

Together, our data suggest that reducing the amount of Shh circulating through the sclerotome promotes a concomitant loss of effective Shh in the NT and vice-versa. Since Hhip:CD4 and Ptc1 are membrane-bound and not secreted, the present results could be explained by the sclerotome, which is a substrate for Shh dispersal, representing a common pool that supplies the ligand to both tissues. Consistent with this notion, sclerotomal Shh is not likely to act by affecting ligand levels in the producing cells because inhibition of Shh by Hhip:CD4 in sclerotome had no effect on Shh expression in either FP or No (Supplem. Fig. 2, E,E’).

### The effects of Hhip:CD4 are a direct consequence of Shh depletion

The similarity between the effects of Hhip:CD4 and Ptc1 (Fig. 2) and the rescue of motoneuron numbers by co-electroporation of Shh along with Hhip:CD4 (Fig. 3) suggests that the effects of Hhip:CD4 are specifically mediated by ligand depletion. To further investigate the possibility of direct versus indirect effects, we examined the expression of *Gli1* and *Hhip1*, two transcriptional targets of Shh and compared it to *Gli3* mRNA expression, which is not directly regulated by Shh (Sasaki et al., 1999). Electroporation of control GFP or of Hhip:CD4 to sclerotome or NT had no effect on expression of *Gli3* mRNA in the same or adjacent tissue (Supplem. Fig. S5, A-C). In contrast, similar transfections of Hhip:CD4 to NT or sclerotome reduced *Gli1* and *Hhip1* mRNAs in both the transfected and in adjacent tissues when compared to the respective contralateral sides (Supplem. Fig. S5, D-I). These results further support the notion that the observed effects on motoneuron and myotome development specifically and directly result from ligand reduction.

Next, we examined the possibility of secondary effects across tissues mediated by Shh depletion. Two candidates are the BMP pathway which acts antagonistically to Shh (Dessaud et al., 2007; Ericson et al., 1997a; McMahon et al., 1998) and retinoic acid from the somite that affects NT development (Diez del Corral et al., 2003; Wilson et al., 2003). To this end, control GFP or Hhip:CD4 were electroporated into the sclerotome. If the effects on the NT of Shh deprivation in sclerotome are mediated by BMP, it is predicted that the dorsal extent of BMP signaling is expanded and or its intensity increased. No changes in the area or intensity of pSmad 1,5,8 immunoreactivity, a readout of BMP activity restricted to the dorsal NT, were measured upon Hhip:CD4 treatment (Supplem. Fig. S6, N=4 per treatment). As an internal control for Hhip:CD4 activity, we observed a reduced number of Hb9+ motoneurons in the ventral NT adjacent to the transfected side (Supplem. Fig. S6B, arrow).

In addition, control GFP or Hhip:CD4 were electroporated into the sclerotome and RARE-AP, a specific reporter of retinoic acid activity was co-electroporated into the NT along with GFP to monitor electroporation efficiency. The specificity of RARE-AP was first tested by co-electroporating it with a dominant negative receptor plasmid, that abolished RARE-AP signal (Supplem. Fig. S7A-B”). No measurable effect in the relative intensity of RARE-AP compared to GFP was observed in the NT upon reduction of Shh in the neighboring sclerotome when compared to control GFP (N= 9 for each treatment, Supplem. Fig. S7, C-E).

Taken together, these results show that the observed effects are a direct consequence of ligand depletion. Thus, sequestering Shh in the sclerotome by Hip:CD4 or Ptc1 results in a corresponding reduction of Shh ligand in the NT and vice-versa.

### Gain of Shh function in sclerotome enhances motoneuron differentiation but Shh missexpression in NT has no effect on myotome

Our loss of function results would be consistent with the possibility that Shh can translocate bidirectionally between mesoderm and NT. To gain additional insight into the directionality of Shh effects across NT and mesoderm, we adopted a complementary gain of function approach. Control GFP or full length Shh were electroporated into sclerotome. A day later, an increase of approximately 30-40% in the number of Hb9+ motoneurons was monitored in the NT of Shh-treated embryos ipsilateral to the treated side when compared to controls (p<0.001, N=9 for each treatment, Fig. 5A-C, see also Fig. 3E).

**Fig.5.**
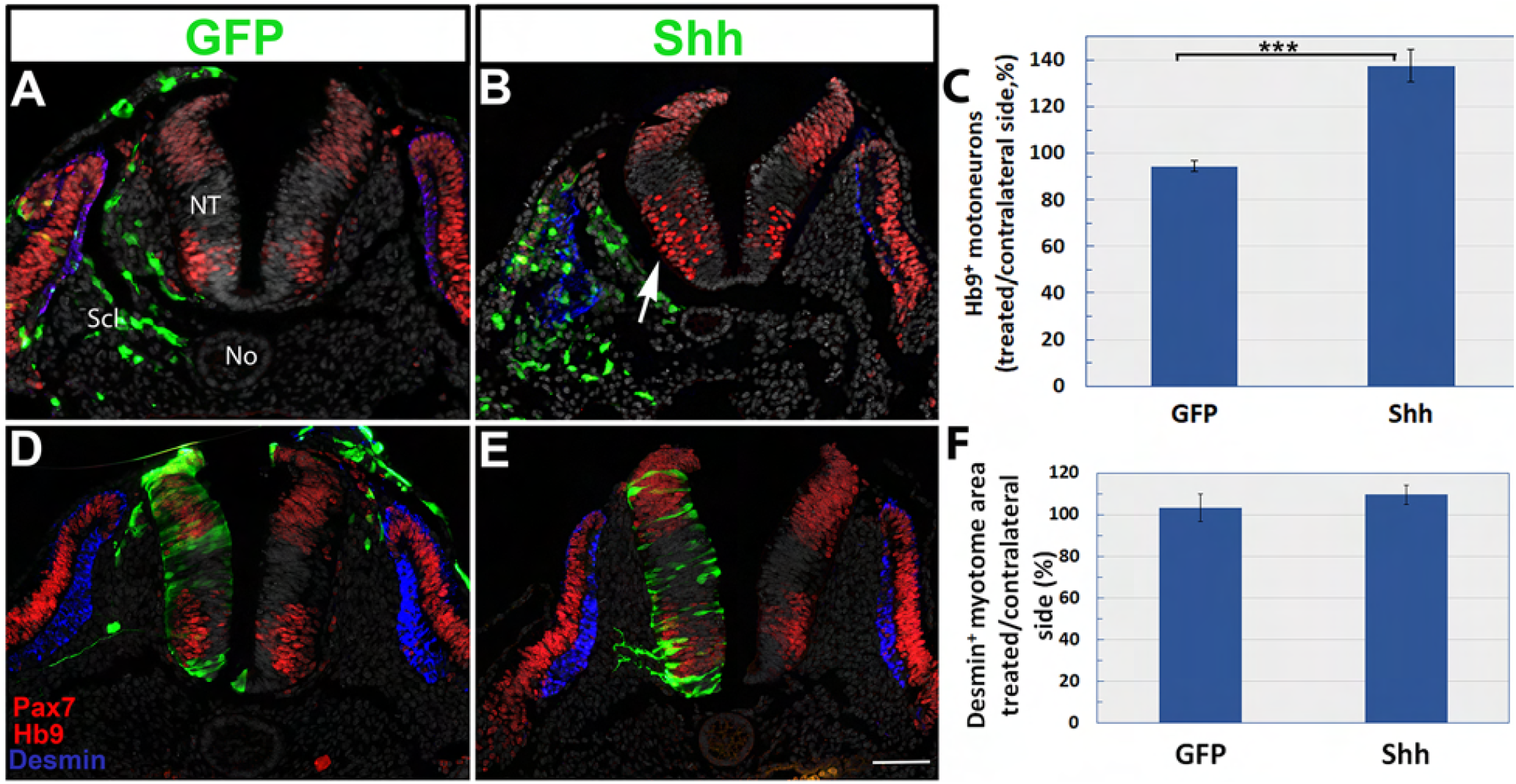
Gain of Shh function in sclerotome enhances motoneuron differentiation but Shh missexpression in NT has no effect on myotome. (A-C) Electroporation of Shh (green) to the sclerotome (Scl) enhances the number of Hb9+ motoneurons in the ventral neural tube (NT) compared to control (arrow in B, red). Quantification in C, ***p<0001. (D-F) Electroporation of Shh (green) to the NT has no effect on the size of desmin+ myotomes (blue) or on expression of Pax7 (red) in the dermomyotme. Quantification in F. Abbreviations, No, notochord; NT, neural tube, Scl, sclerotome. Bar=50 mM.

Reciprocally, control GFP or full length Shh were electroporated into hemi-NTs and the size of adjacent desmin+ myotomes was examined. Although the overall size of the transfected hemi-NT increased, no significant change in myotomal size was measured and no apparent effects on DM or sclerotome were observed (N=6 and 5 in control and treated embryos, respectively, Fig. 5D-F).

Hence, our gain of function data are inconsistent with the simpler possibility that Shh moves bidirectionally between mesoderm and NT, that is based solely on data from loss of Shh activity. While excess Shh in mesoderm profoundly affects NT development, the observation that gain of Shh in NT has no effect on myotome development is in line with Shh being transported into the neuroepithelium but not outside into mesoderm. In this regard, our finding that Shh depletion in the NT inhibits myotome differentiation (Fig. 4) is consistent with this procedure causing a corresponding lower effective concentration in mesoderm. This could be accounted for by enhanced uptake of the ligand into NT cells via directional baso-apical transport from the sclerotome. This suggests that the sclerotome constitutes a common pool of No-derived Shh that serves both tissues.

### Sclerotome, but not FP-derived Shh is necessary for motoneuron differentiation

The currently accepted view sustains that the development of various cell types in the NT depends upon a local ventrodorsal gradient of Shh directly emanating from the No and/or FP (Smith, 1993; Trousse et al., 1995). Our present results, show that the sclerotomal pool of Shh, originally released from No/FP, is also necessary for motoneuron development (Figs. 4,5). Hence, we examined the relative contribution of the FP compared to sclerotomal Shh to the differentiation of Hb9+ neurons.

Control GFP or Hhip:CD4 were electroporated dorsoventrally to attain the FP of the NT. Control GFP had no effect on FP integrity or on expression of Shh protein (N= 7, Fig.6A). Surprisingly, electroporation of Hhip:CD4 caused by 24 hours, the total absence of the FP and consequent loss of FP-derived Shh (N= 10, Fig. 6, D,E). Notably, already by 10 hours post-electroporation, the beginning of FP disintegration, reflected by the presence of pyknotic nuclei, loss of Shh and punctate appearance of transfected Hhip:CD4-GFP were observed (Fig. 6G-G”, arrows and arrowheads, N=5). This early loss of FP enabled us to accurately monitor the contribution of the FP to motoneuron development. Because the effect is bilateral, in order to prevent variability between embryos and obtain a reliable measurement of motoneuron numbers, the proportion of Hb9+ neurons was measured as the ratio between the neurons present at the flank level lacking a FP to the intact neck level of the same embryos. In spite of FP disappearance, the proportion of flank-level motoneurons was unaltered when compared to control embryos that received GFP only (N=4 for each treatment, Fig. 6, B,E,I). Similar to what was observed with Hhip:CD4, dorsoventral electroporation of Ptc1 also compromised FP integrity while having no apparent effect on motoneurons (N=5, Fig.6H). In contrast, inhibition of Shh in the sclerotome by Hhip:CD4 exhibited a visible reduction in ventral motoneurons (Fig. 6, C,F arrow, and see also Figs. 2 and 3). Thus, we conclude that Shh traversing the sclerotome plays a significant part in motoneuron differentiation whereas the FP itself has no apparent contribution at least at the stages examined in our study (see Discussion).

**Fig. 6.**
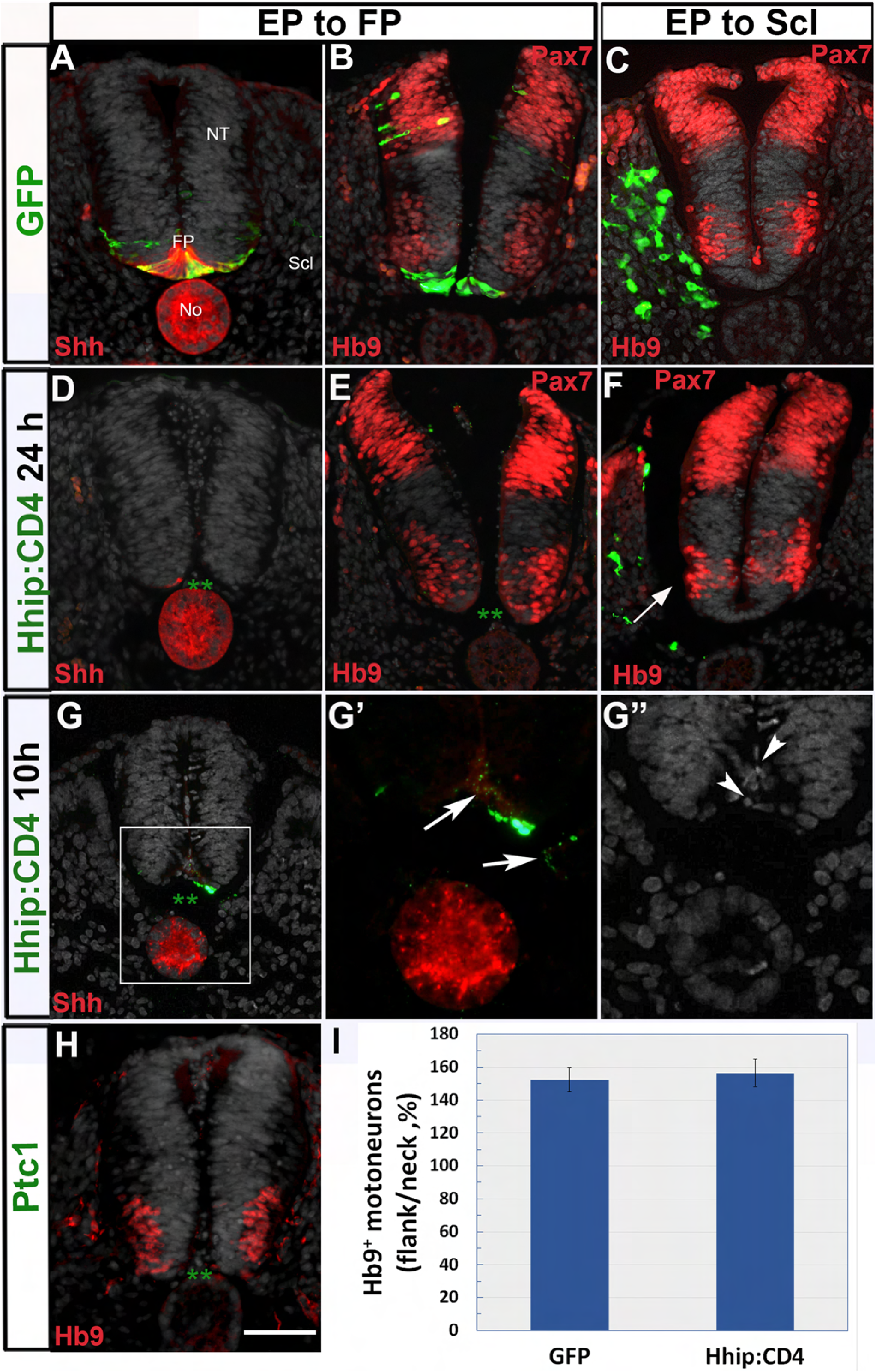
Sclerotome, but not FP-derived Shh is necessary for motoneuron differentiation. (A,B) Dorsoventral electroporation of control GFP showing in A, the presence of the labeled floor plate (FP) that co-expresses Shh protein. In B, note Hb9+ motoneurons dorsal to the labeled FP. (C) Hb9+ motoneurons in control GFP electroporation to sclerotome. (D,E) Loss of FP tissue 24 hours post-Hhip:CD4 electroporation. Asterisks (**) in D and E denote absence of a FP and panel D shows concomitant absence of Shh expression. Hhip:CD4 electroporation has no effect on ventral Hb9+ motoneurons or dorsal Pax7 expression (both in red). (F) In contrast, less motoneurons are apparent adjacent to the sclerotome transfected with Hhip:CD4 (arrow). (G-G”) Disintegration of the FP 10 hours following Hhip:CD4 transfection. Note in G and G’ the absence of Shh immunoreactivity in the FP domain as well as punctate expression of GFP (arrows). In G” note the presence of disorganized and pyknotic nuclei. (H) Loss of the FP upon electroporation of Ptc1 (**) shows no apparent effect on motoneurons. (I) Quantification showing no effect of FP deletion on the proportion of Hb9+ motoneurons in flank (electroporated)/neck (intact region). Abbreviations, No, notochord; NT, neural tube. Bar=50 μM.

### A basal, but not apical, presentation of Shh is required for ligand activity on both NT and DM/myotome

Based on our finding that Shh transiting through the sclerotome is needed for NT development, we predicted that the NT would be more sensitive to Shh presented from its basal side abutting the sclerotome than from its apical side. To examine this hypothesis, fragments of No were grafted into either the lumen of the NT (apical) or between somites and NT (basal grafts). A day later, the there was no apparent reduction in the extent of Pax7 expression in the luminal grafts; and only a mild reduction in the intensity of Pax7 was monitored (Fig. 7A, compare with control side of panel B). In contrast, Pax7 immunoreactivity was dorsally restricted when facing the basal grafts (Fig. 7B). The latter also caused a characteristic bending of the NT, previously reported to represent an ectopic FP-like structure (van Straaten et al., 1988)(N= 6 and 6 for apical vs. basal grafts, Fig. 7, B,D). In addition, the amount of Hb9+ motoneurons was unchanged by the apical grafts, yet was markedly increased in the basal grafts in which the No’s were similarly localized in a dorsal position vis-vis the NT (N= 6 and 6 for apical vs. basal grafts, Fig. 7, C,D), in agreement with classical No graft experiments (van Straaten et al., 1989).

**Fig. 7.**
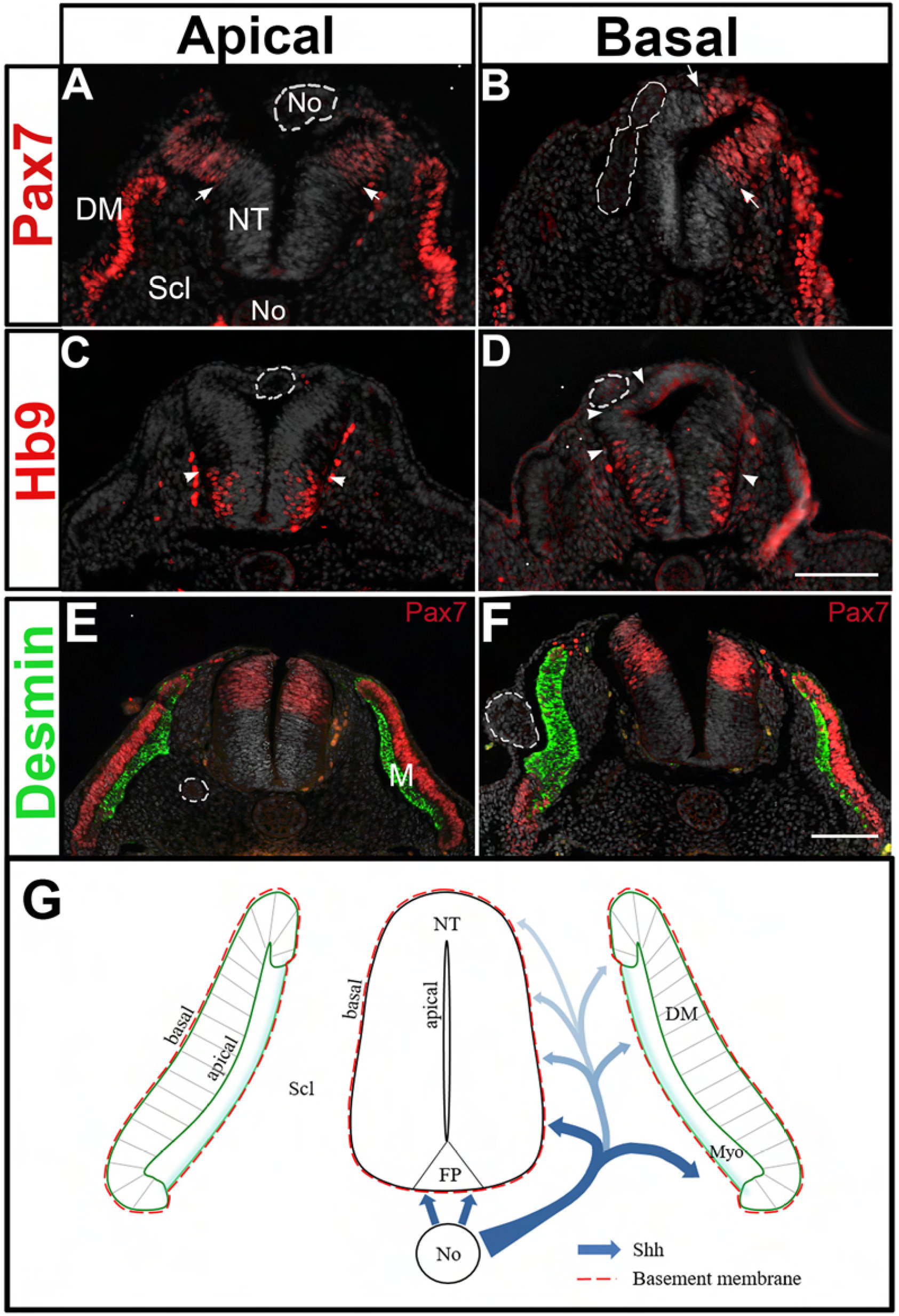
A basal, but not apical, presentation of Shh is required for ligand activity on both NT and DM/myotome. A model accounting for the effects of Shh traversing the sclerotome on both NT and myotome development. Implantation of No fragments (dotted circles) in apical (A,C,E) or basal (B,D,F) positions vis-à-vis the dorsal NT, or the dermomyotome (DM) (E,F). (A,C) A No piece was grafted inside the NT facing its luminal side. Note that a day later it localizes in the dorsal NT. No effect on Pax7 (arrows) or Hb9+ motoneurons (arrowheads) is apparent. (B,D) A No piece was grafted in a basal position vis-à-vis the dorsal NT. Note a dorsal restriction of Pax7 (arrows in B) as well as a dorsalward expansion of Hb9+ motoneurons (arrowheads in D). (E,F) A No fragment localized apical to the DM causes a mild change in the adjacent desmin+ myotome (green, E) whereas the basally located No strongly increases myotomal size and promotes an in situ differentiation of the DM into myotome with concomitant loss of Pax7 expression. (G) A proposed model for the activity of No-derived Shh. The No secretes Shh that acts on the ventral NT and also traverses the sclerotome (Scl) which is both a pathway for ligand movement and also a target of its activity. We propose that a ventral to dorsal gradient of ligand is created in Scl and plays a pivotal role both on myotome differentiation as well as on motoneuron development. Shh is thus presented to the target epithelial cells via its basal domain, probably by initial association with the laminin-containing basement membrane. See text for details. Bar=50 μM.

Likewise, implantation of No fragments at epithelial somite levels and at an apical position with respect to the DM had only a mild effect on the subsequent development of desmin-positive myotomes with no apparent alteration in Pax7 expression in the DM. In striking contrast, equivalent grafts performed basal to the prospective DM, produced large myotomes expressing desmin and a radical *in situ* differentiation of the DM into muscle at the expense of Pax7-positive progenitors (N= 5 and 7 for apical vs. basal grafts, Fig. 7, E,F). Therefore, an initial basal presentation of the ligand vis-à-vis the target epithelium is required for the activity of Shh. This is consistent with our results showing that Shh emanating from the sclerotome and reaching the NT from its basal side, is necessary and sufficient for aspects of NT differentiation.

## Discussion

In the present study we uncover a previously unknown domain, the sclerotome, as being an important “en passant” substrate of Shh that influences not only DM and myotome development, as previously shown (Kahane et al., 2013) but also aspects of NT differentiation (Fig. 7G). Our loss of function data show that local depletion of Shh in either NT or sclerotome results in major defects across tissues such as reduced myotomal size and less motoneurons, respectively. Reciprocally, only gain of Shh function in the sclerotome significantly enhances motoneuron differentiation while missexpression of the ligand in NT has no effect on myotomal size. Taken together, these results suggest that Shh ligand specifically secreted into in sclerotome constitutes a common pool that supplies both tissues.

In contrast to the reduction in motoneurons observed upon inhibition of Shh in sclerotome, we show that Hhip:CD4-mediated ablation of the FP has no short-term effects on motoneuron numbers. This is consistent with the development of a normal ventral pattern in Gli2 mutants that lack a FP (Chamberlain et al., 2008; Matise et al., 1998a). Likewise, loss of Shh in FP did not dramatically alter ventral neural patterning, yet altered gliogenesis at a later stage (Yu et al., 2013). Furthermore, abrogating Shh in FP under the regulation of Brn4, revealed a normal short-term expression of both *Nkx2*.*2* and *Olig2*, but a reduction at later stages suggesting a continuous need for FP-derived Shh in development of the ventral NT (Dessaud et al., 2010). This initial phenotype is consistent with our results yet we did not analyze later stages. The above results indicate that Shh required at this stage for motoneuron development does not derive from the FP, which is an integral part of the neuroepithelium and from which the ligand could potentially disperse apically or through the cells themselves. It is therefore possible that in the absence of a FP, the No compensates for the loss of FP-derived Shh as suggested for Gli2 mutants [reviewed in (Danesin and Soula, 2017)]. In such a case, No-derived Shh would follow a basal pathway. Consistently, abrogation of Shh ligand in mesoderm, performed at the same stage and for a similar duration, revealed a significant NT phenotype even in the presence of both Shh-producing axial structures. Taken together, we propose that at least a significant fraction of Shh operating on the neuroepithelium stems from the No via a sclerotomal pathway which, at the stages examined, seems more active than the FP in promoting motoneuron development. Along this line, we could not address effects of Shh reduction in the earlier mesoderm on initial ventrodorsal NT patterning because of technical limitations of the electroporation procedure. Instead, we monitored later events (e.g; cell proliferation, survival and differentiation), all known to depend upon Shh. Thus, the possibility remains open that Shh circulating through mesoderm contributes to NT development at numerous stages.

Furthermore, if Shh from sclerotome is important for neuroepithelial development, is it possible to directly show its presence in this domain? Shh in the synthesizing cells of the FP and No is intracellular and membrane-bound [(Chamberlain et al., 2008) and Supplem. Fig. S2)] whereas in the sclerotome it is expected to be extracellular and/or included in organelles (e.g, exosomes, etc). Most protocols used for tissue processing are likely to keep only the former type of immunoreactive protein and wash away the extracellular ligand in sclerotome. Notably, when using a method that allowed proteoglycan/glycosaminoglycan preservation and/or perhaps also a different antibody, a previous study showed a sclerotomal localization of Shh immunoreactive protein (Gritli-Linde et al., 2001).

It is noteworthy that different procedures that perturb the production of Shh protein or its signaling have different effects on FP integrity. Whereas Gli2 mutants lack a FP (Matise et al., 1998b), deletion of Shh in FP does not affect maintenance of this structure (Dessaud et al., 2010). In both chick and mouse, Shh was suggested to be necessary for initial FP induction but later on, during somitogenesis, the FP becomes refractory to the ligand (Ribes et al., 2010). In our experiments, we implemented a membrane tethered version of the high affinity Shh inhibitor Hhip and also the Shh receptor Ptc1, both resulting in the death of FP cells. This might be accounted for by a combination of ligand depletion with accumulation of Shh-Hhip or Shh-Ptc1 complexes at the cell membrane altogether compromising the structural integrity of this epithelium.

One open question stemming from our results is how is Shh transported through the sclerotome. Possible models could be packaging of the ligand in No-derived exosomes (Vyas et al., 2014), diffusion of Shh released by matrix metalloproteinases in a lipid-free form (Dierker et al., 2009); secretion of Shh as multimeric complexes of various molecular compositions [see for example (Chen et al., 2004)] and/or via carrier-mediated transport through the extracellular space (Parchure et al., 2018). The precise mechanism responsible for Shh transport in the present context remains to be unraveled.

If provided to the NT from the sclerotomal domain, it is inferred that neurepithelial cells sense Shh from their basal pole that faces the mesoderm. Consistent with this notion, grafting No fragments in a basal, but not apical position with respect to the NT, profoundly affects NT shape, motoneuron differentiation and Pax7 expression. Indeed, during normal development, the No underlies the basal domain of the NT, further supporting the idea that No-derived Shh can only reach the NT via a basal route. In line with the above, it was reported that lipidated Shh enters the cells of the imaginal disc in Drosophila only through its basolateral surface (Callejo et al., 2006). A similar phenomenon was observed in high density human gastruloids, that self organize into an epithelium. In these cultures, cells were responsive to BMP4 or Activin ligands only when presented from the basal side and this correlated with the localization of BMP receptors at the basolateral domain of the cells (Etoc et al., 2016). Although it is unclear if Shh receptors are also localized at the basal aspect of neuroepithelial cells, or if Shh needs first to be transported baso-apically in order to act on apically localized receptors, all the above results would suggest that cell polarization controls ligand response.

The above findings are interesting in light of the proposed concept that apically-localized cilia serve as antennae to sense and transduce a Shh signal (Corbit et al., 2005; Singla and Reiter, 2006). Based on our data, we suggest that, initially, Shh is presented from the basal side of epithelial cells from which it may be transported to the apical domain where cilia are localized. Since an apical presentation of the No and associated Shh was without a significant effect in our implant experiments, we propose that cilia act primarily as transducers of a Shh signal, but not as the primary antennae sensing the presence of the ligand. Moreover, the observation that a direct apical presentation of Shh is without a major effect, further suggests that basal reception followed by baso-apical transport might be necessary for the activity of Shh arriving at the cilia. Along this line, growing evidence suggests cilia-independent Shh reception that occurs through basally localized cytonemes [reviewed in (Gradilla et al., 2018) and refs therein]. Likewise, in the retina neuroepithelium, Shh and its coreceptor Cdo colocalize at the basal side of the cells where filopodia-like structures are present (Cardozo et al., 2014). An extreme example is provided by some cell types where Shh signaling takes place even in the absence of cilia [reviewed in (Gradilla et al., 2018) and refs. therein].

In the NT, direct visualization of the behavior of labeled Shh, revealed that the ligand from the No concentrates in association with the apically localized basal bodies from which cilia stem, while forming a dynamic gradient in the ventral NT (Chamberlain et al., 2008). In light of the present results, the above observed graded distribution of ligand could be explained as being the end point of a transport process that begins at the No, travels through the sclerotome forming there a ventro-dorsal gradient, then reaches the neuroepithelium through its basal side finally concentrating in the apical cilia (Fig. 7G). In such a model, the mechanisms mediating the baso-apical transport of the ligand associated with possible biochemical changes of the protein that make it availabile to cilia for signaling remain to be unraveled. A microtubular network spanning the extent of neuroepithelial cells could be involved in this process, as previously suggested (Chamberlain et al., 2008).

An initial presentation of Shh from the basal side of an epithelium seems to be of general significance as basal grafting of a No also elicited robust *in situ* differentiation of DM progenitors into myocytes when compared to an apical implant that exhibited only a mild phenotype. How can this differential effect be explained given that the endogenous ligand apparently arrives from the apical sclerotomal direction? Initially, pioneer myotomal cells in the epithelial somite face the No from their basal aspect (Kahane et al., 1998). With ongoing development and upon sclerotome dissociation, the epithelial DM is consolidated and becomes composed of a central sheet and four inwardly curved lips pointing towards the sclerotome. In previous studies, we and others showed that the four lips of the DM are the main sources of myotomal cells [(Kalcheim et al., 1999) and refs therein] and it is their basal domain that points towards the sclerotome from which Shh arrives. Next, myotomal progenitors enter the nascent myotome and a basement membrane begins assembling at the interface between myotome and sclerotome (Borycki, 2013). At this time, the central DM sheet also contributes myotomal progenitors by direct translocation into the myotome (Ben-Yair et al., 2011), which then differentiate in a Shh-dependent manner (Kahane et al., 2013). These precursors were shown to exhibit an inverse apicobasal polarity when compared to the central DM cells (Ben-Yair et al., 2011). Hence, both the DM lips and these translocating progenitors point with their basal surfaces towards the source of endogenous Shh, likely being the main targets for its activity and also accounting for the partial effect of the apical grafts.

As discussed above, the basal domain of the DM/myotome and NT epithelia are characterized by the presence of a surrounding basement membrane. An association between Shh and the basal lamina has been shown. For instance, in cerebellar granule cell precursors, the laminin-containing basement membrane binds and locally enhances Shh signaling (Blaess et al., 2004). Similarly, in the mouse somite, Shh induces the activation of Myf5 in DM. Myf5+ cells then translocate to the myotome and upregulate α6β1 integrin and dystroglycan, allowing a myotomal basement membrane to be assembled using primarily laminin α1 produced by sclerotomal cells, and laminin α5 produced by the dorsomedial lip of the DM [reviewed in (Borycki, 2013)]. This is further confirmed in Shh-deficient mice, which fail to form a myotomal basement membrane, and in which myotomal cells do not properly exit the cell cycle, maintain Pax3 expression and delay the differentiation program (Anderson et al., 2009; Borycki et al., 1999). Additionally, Shh immunoreactive protein was found to localize in the basement membrane surrounding the NT (Gritli-Linde et al., 2001). Taken together, these results suggest that in both NT and DM/myotome, a feedforward mechanism may exist whereby Shh controls laminin expression and the assembly of a basement membrane. This could allow a local concentration of the ligand and/or signal stabilization. However, the basement membrane alone is unlikely to serve as the common pool, as sequestering the ligand specifically in sclerotome by membrane-associated Hhip:CD4 was highly effective. In addition, others have shown that the ligand is present in the mesoderm itself (Gritli-Linde et al., 2001).

Our present findings provide an additional argument in support of the importance of NT-somite interactions which are pivotal for the normal patterning of trunk components. For instance, opposite gradients of retinoic acid and Fgf8 in mesoderm are required for early NT development (Diez del Corral et al., 2003). In addition, the nascent DM controls the timing of neural crest delamination by modulating noggin mRNA and BMP activity levels in the NT (Sela-Donenfeld and Kalcheim, 2000). Reciprocally, Bmp4 and/or Wnt1 from the dorsal NT pattern the somite-derived DM and affect myogenesis (Abu-Elmagd et al., 2010; Marcelle et al., 1997). Formation of the dorsal dermis from the DM is influenced by NT-derived Wnt1 and by neurotrophin-3 (Brill et al., 1995; Marcelle et al., 1997; Sela-Donenfeld and Kalcheim, 2002). Furthermore, interactions between neural and somitic cells control neural crest migration and segmentation of peripheral ganglia and nerves as well as specific aspects of myogenesis (Kalcheim, 2011; Kalcheim and Goldstein, 1991). The present results raise the intriguing possibility that the dual activity on both motoneurons and myotome of Shh released into sclerotome, serves to couple and coordinate development of the neuromuscular system.

## Materials and Methods

### Embryos

Chick (Gallus gallus) and quail (Coturnix japonica) eggs were from commercial sources (Moshav Orot and Moshav Mata, respectively). All experiments were performed using quails except for those involving in situ hybridizations which were done in chick embryos.

### Expression vectors and electroporation

Expression vectors were: pCAGGS-GFP, pCAGGS-RFP (Krispin et al., 2010), Ptc1 (Briscoe et al., 2001), PTC^Δloop2^ (Briscoe et al., 2001; Kahane et al., 2013), a retinoic acid reporter fused to alkaline phosphatase (pRARE-AP, from J. Sen) (Gupta and Sen, 2015), a dominant negative pan-retinoic acid receptor (RAR403dn-IRES-GFP, from S. Sockanathan) that abrogates retinoic acid signaling (Novitch et al., 2003), full length Shh (Kahane et al., 2013), cholesterol deficient Shh (mShh-N-YFP, from V. Wallace) (Beug et al., 2011) that was subcloned into pCAGGS, and mHhip1 (Kahane et al., 2013). To produce Hhip1:CD4, a membrane-tethered version of Hhip1, the transmembrane and intracellular domains of mouse CD4 were fused to the C-terminal domain of Hhip1 lacking amino acids A679-V700, as previously described (Holtz et al., 2015; Kwong et al., 2014) and further subcloned into pCAGGS for electroporation.

For electroporations, DNA (1-4 µg/µl) was microinjected into the center of flank-level epithelial somites (somites 20-25) of 23-25 somite-stage embryos. Electroporations were performed to the ventral half of epithelial somites (prospective sclerotome). To this end, the positive tungsten electrode was placed under the blastoderm in a location corresponding to the ventro-medial portion of epithelial somites on a length of about 7 segments, and the negative electrode was placed in a superficial, dorso-lateral position with respect to the same somites. (Ben-Yair et al., 2011; Halperin-Barlev and Kalcheim, 2011; Kahane et al., 2007; Kahane et al., 2013). For hemi-NT electroporations, DNA was microinjected into the lumen of the NT. One tungsten electrode was placed underneath the blastoderm on one side of the embryo and the other electrode was placed in a superficial position at the contralateral side. For FP electroporations, the positive electrode was inserted under the blastoderm near the midline and the negative electrode was placed over the dorsal NT. In some cases double electroporations to the hemi NT and sclerotome were performed sequentially. A square wave electroporator (ECM 830, BTX, Inc.) was used. One pulse x12V for 5msec was applied.

### No grafts

No fragments comprising a length of 7-8 segments were enzymatically excised from donor embryos aged 25 somite pairs as previously described (Charrier et al., 2001) and kept in cold phosphate buffered saline until grafting. For grafting at the apical side of the NT, the ectoderm and dorsal NT of host embryos were cut and the No piece placed in the NT lumen. A day following implantation, the grafts were usually found in the dorsal portion of the NT facing its cavity. For basal grafting with respect to the NT, a slit was performed between somites and NT and the No fragments were inserted. A similar, but more profound slit, was performed to reach the ventral sclerotome abutting the apical side of the DM. To reach the basal domain of the DM, the ectoderm was cut to precisely accommodate the length of the No fragment. Following microsurgery, embryos were reincubated for additional 24 hours.

### Immunohistochemistry

Embryos were fixed overnight at 4°C with 4% formaldehyde in phosphate-buffered saline (PBS) (pH 7.4) followed by washings in PBS. Most immunostainings except for desmin were performed on whole embryo fragments. Immunolabeling for desmin was performed on tissue sections, as described (Burstyn-Cohen and Kalcheim, 2002; Kahane et al., 2001).

For wholemount immunostaining, antibodies were diluted in PBS containing 1% Triton X-100 and 5% newborn calf serum and tissues were incubated overnight at 4°C on a rotatory shaker. Next, they were washed twice in a large volume of PBS/1% Triton X-100 first for 10 min and then for 2 hours at room temperature. Secondary antibodies were similarly diluted in PBS/ 1% Triton X-100/ 5% newborn calf serum and incubated overnight followed by repetitive washings. Embryo fragments were dehydrated in increasing ethanol solutions (30%,70%, 90% and 100%, 10 minutes each) followed by toluene (2 times, 10 minutes each), then embedded in paraffin wax and sectioned at 8μm. Paraffin was removed in Xylene and slides were rehydrated in decreasing ethanol solutions.

The following antibodies were used: rabbit anti GFP (1:2000, Invitrogen, Thermo Fisher Scientific, A6455) and mouse anti-desmin (1:200, Molecular Probes, 10519). Monoclonal antibodies against Pax7 (PAX7-s, 1:20), Shh (5E1, 1:20) and Hb9 (1:200) were from DSHB, University of Iowa). Phosphorylated Smad 1-5-8 (PSmad, 1:1000) was a gift from Ed Laufer. Anti-Histone H3 (phospho S10) was from Abcam (mAbcam 14955, 1:500). Detection of DNA fragmentation was done by TUNEL (ab66110, Abcam) according to manufacturer’s instructions. Nuclei were visualized with Hoechst.

### In situ hybridization

Embryos were fixed in Fornoy (60% ethanol, 30% formaldehyde, 10% acetic acid), then dehydrated in ethanol/toluene, processed for paraffin wax embedding and sectioned at 10 μm. Slides were rehydrated in toluene/ethanol/PBS, treated with proteinase K (1µg/ml, Sigma Aldrich P2308) at 37°C for 7 minutes, and then fixed in 4% formaldehyde at room temperature for 20 minutes. Next, slides were washed in PBS followed by 2X SSC and hybridized in hybridization buffer (1X salt solution composed of 2M NaCl, 0.12M Tris, 0.04M NaH_2_PO_4_2H2O, 0.05M Na_2_HPO_4_, 0.05M EDTA, pH7.5], 50% formamide, 10% dextran sulfate, 1mg/ml Yeast RNA, 1X Denhardt solution) containing 1μg/ml DIG labeled RNA probes (prepared with a DIG RNA labeling mix, Roche, 11277073910) for overnight at 65°C in a humid chamber. Post-hybridization, slides were rinsed in a rotating incubator with 50% formamide, 1X SSC, 0.1% Tween 20, until coverslips dropped and then an additional wash for 1 hour followed by 2 washes in MABT (10% Maleic acid 1M pH 7.5, 3% NaCl 5M, 0.1% Tween 20) and preincubation in MABT/ 2.5% FCS. Anti-DIG-AP antibody (1/1000, Roche 11093274910) diluted in MABT+2% BBR+20% FCS was then added for overnight at room temperature. This was followed by rinsing in MABT and then in NTMT (2% NaCl 5M, 10% Tris HCl 1M pH9.5, 5% MgCl_2_ 1M, 0.1% Tween20), and then incubation in NTMT + 1:200 NBT/BCIP Stock Solution (Sigma-Aldrich, 11681451001) at 37°C until the AP reaction was completed.

The following probes were employed: *Hhip1* (from J. Briscoe) *, Nkx*2.2, *Nkx* 6.1, *Nkx* 6.2, and *Olig2* (from J. Ericson), and *Gli1*, *Gli3* from A.G. Borycki.

### Data analysis and statistics

Four to 26 embryos were analyzed per experimental treatment. The number of Hb9-positive motoneurons or was counted in 5-10 alternate sections per embryo. The number of phospho-histone H3 (pH3) + nuclei was monitored in 7-10 sections per embryo in 5-8 embryos per treatment.

Myotomes were defined by desmin staining. The area occupied by desmin+ myotomes was measured in alternate sections of 3 to 16 embryos per experimental treatment. Sclerotomes of 4 to 6 embryos were defined in the mediolateral aspect as the tissue between myotome and NT, and in the dorsoventral extent as the mesenchyme between the dorsomedial lip of the DM up to the dorsal border of the cardinal vein and aorta. The surface area of hemi-NTs was monitored in 4 sections per embryo. Myotomal, sclerotomal and hemi-NT areas as well as the area and intensity of cells expressing the p-Smad 1,5,8 or pRARE-AP were measured using Image J software (NIH). All results are expressed as the mean proportion of positive cells or area in treated compared to control contralateral sides, or to control GFP ±SEM.

In a control experiment, we tested whether area measurements are a faithful representation of cell number. To this end, we compared the number of Hoechst+ nuclei in hemi-NTs electroporated with Hhip:CD4/contralateral side vs. control GFP/contralateral side, and found them to be not significantly different from the equivalent ratio of surface area. Consequently, the calculated ratio of area to cell number was similar (1.03±0.032, N=6 embryos sectioned at 5μM, 5 sections/embryo). Thus, surface area was validated as a reliable measure of overall cell number.

Images were photographed using a DP73 (Olympus) cooled CCD digital camera mounted on a BX51 microscope (Olympus) with Uplan FL-N 20x/0.5 and 40x/0.75 dry objectives (Olympus). For quantification, images of control and treated sections were photographed under the same conditions. For figure preparation, images were exported into Photoshop CS6 (Adobe). If necessary, the levels of brightness and contrast were adjusted to the entire image and images were cropped without color correction adjustments or γ adjustments. Final figures were prepared using Photoshop CS6.

Significance of results was determined using the non-parametric Mann–Whitney test. All tests applied were two-tailed, and a *P*-value of 0.05 or less was considered statistically significant. Data were analyzed using the IBM SPSS software version 25. The number of embryos analyzed for each treatment (N) is detailed in the Results Section. P-values can be found both in the Results Section and in the corresponding Legends.

## Acknowledgements

We thank Mordechai Applebaum and Dina Rekler for help with figure preparation and Tali Bdolach for assistance with statistics. We thank James Briscoe and Johan Ericson for helpful comments during the course of this study, and Joel Yisraeli for critical reading of the manuscript. This work was supported by the Israel Science Foundation (#97/13) to CK.

## Author Contributions

NK and CK conceived the project and designed the experiments; NK performed the experiments; CK and NK wrote the paper.

## Data availability

Not applicable.

## Legends to Supplementary Figures

**Supplementary. Fig. S1.**
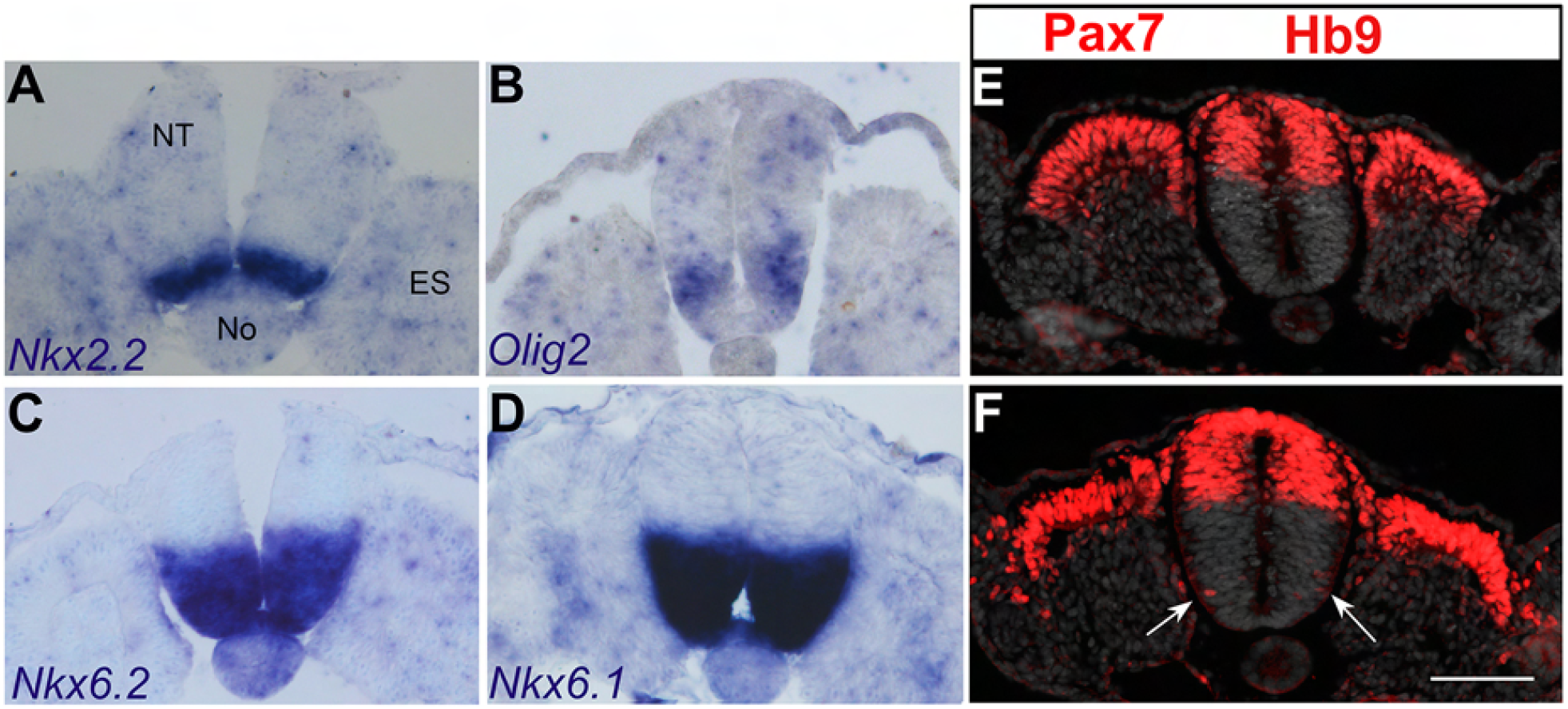
Expression of neural tube markers by the time of *in ovo* electroporation. (A-D) In situ hybridization of embryos aged 25 somite pairs showing expression of ventral neural tube (NT) markers at the epithelial somite (ES) stage. (E,F) Co-immunolabeling of Pax7 and Hb9 antibodies. Note in E that only Pax7 is expressed in both dorsal NT and dorsal aspect of the ES, whereas the first Hb9+ ventral motoneurons appear at a later stage following somite dissociation (arrows in F). Both dorsal Pax7 and ventral Hb9 immunostainings are in red as the respective antibodies are monoclonal of the same subtype. Abbreviations, No, notochord. Bar=50μM.

**Supplementary Fig. S2.**
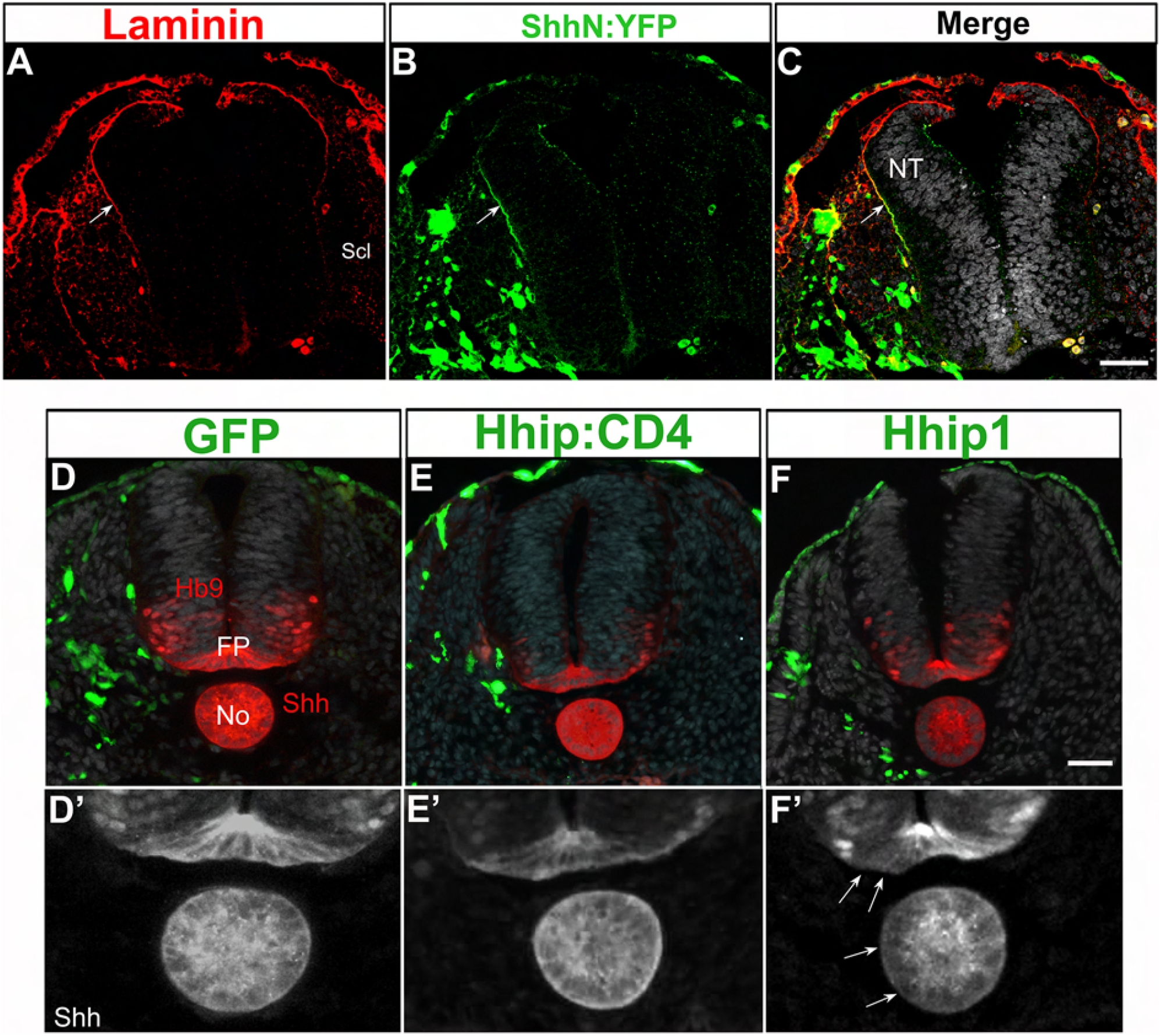
Differential behavior of secreted Hhip1 compared to membrane-tethered Hhip:CD4. (A-C) ShhN:YFP, electroporated into sclerotome (Scl, green), colocalizes with laminin (red) in the basement membrane around the neural tube (NT) (arrows). (D-F’) Immunolabeling with Shh and Hb9 antibodies of embryos electroporated with control GFP (D,D’), Hhip:CD4 (E,E’) or Hhip1 (F,F’). Note that Hhip1 caused a reduction in ipsilateral Shh in both notochord (No) and FP (FP) (arrows in F’) whereas GFP or Hhip:CD4 were without effect. Bars=50 μM.

**Supplementary. Fig.S3.**
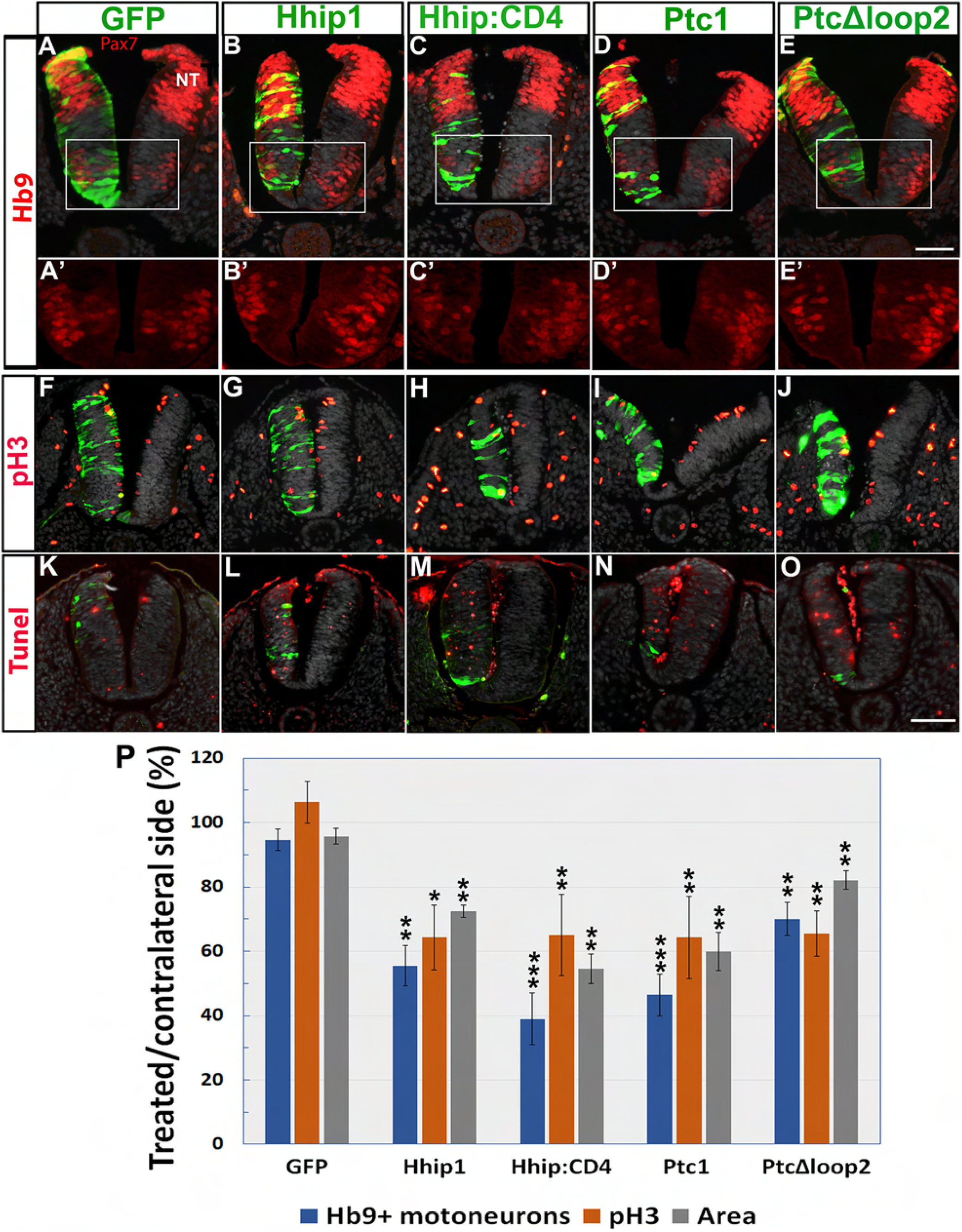
Depletion of Shh in the NT by Hhip:CD4 phenocopy those exerted by Hhip, Ptc1 and PTC^Δloop2^. (A-E’) Electroporation of depicted plasmids (green) to hemi-NTs followed a day later by immunostaining for Pax7 and Hb9. A’-E’ represent higher magnifications of the boxed regions in A-E. (F-J) staining of mitotic nuclei with anti pH3 (red) following electroporation with depicted plasmids. (K-O) Tunel staining (red). Note weak immunostaining of electroporated plasmids (green) because no anti-GFP antibodies were implemented to enable better visualization of apoptotic nuclei. (P) Quantification of motoneurons, mitotic nuclei and area. *p<0.05, **p<0.03, ***p<0.01. Bar=50μM.

**Supplementary Figure S4.**
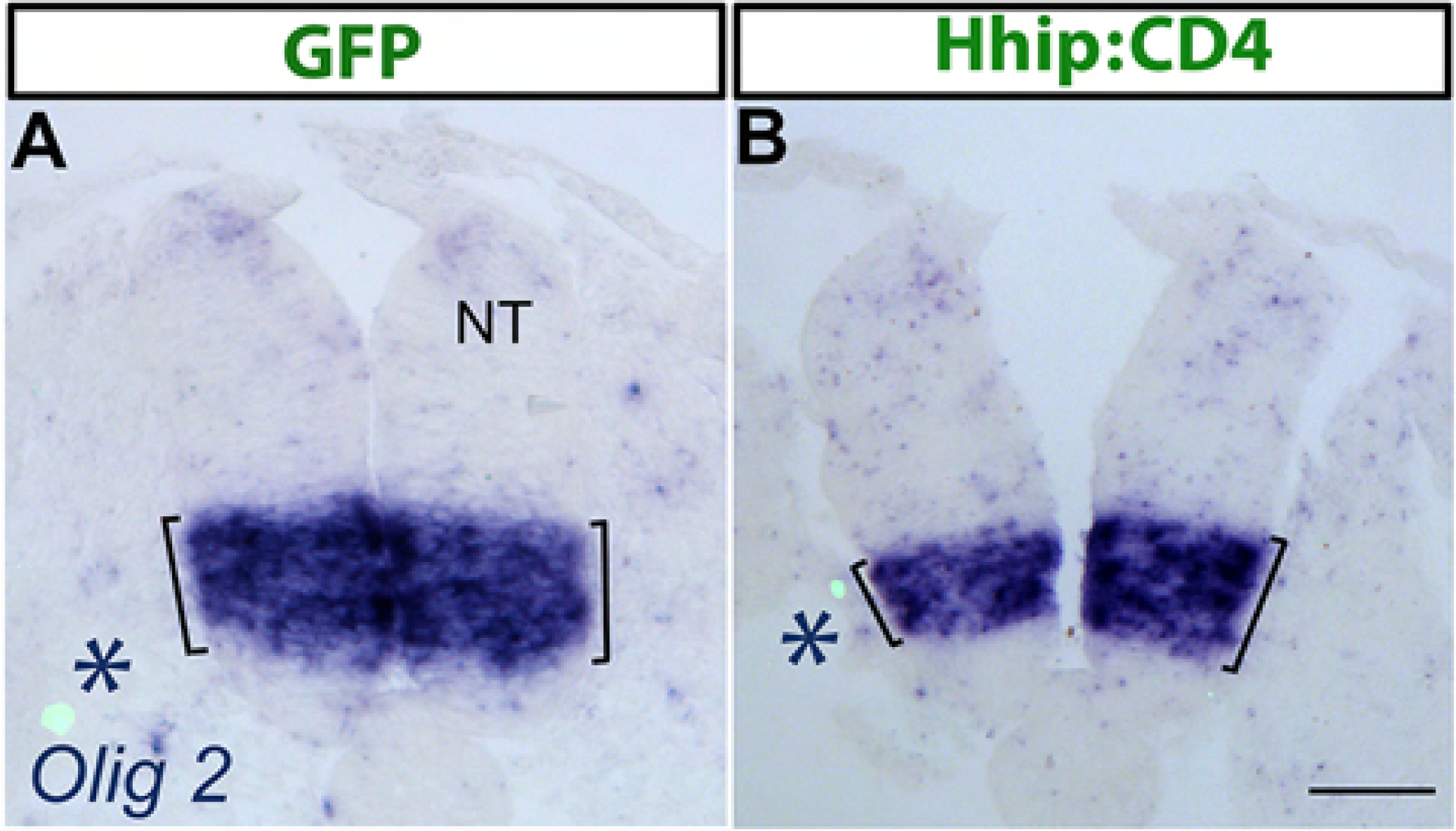
Electroporation of Hhip:CD4 into the sclerotome reduces the extent of *Olig2* expression in NT. (A,B) Electroporation of control GFP or Hhip:CD4, respectively. Asterisks denote the transfected sclerotomes. The extent of *Olig2* mRNA expression is delimited by black lines. Bar=50μM.

**Supplementary. Fig. S5.**
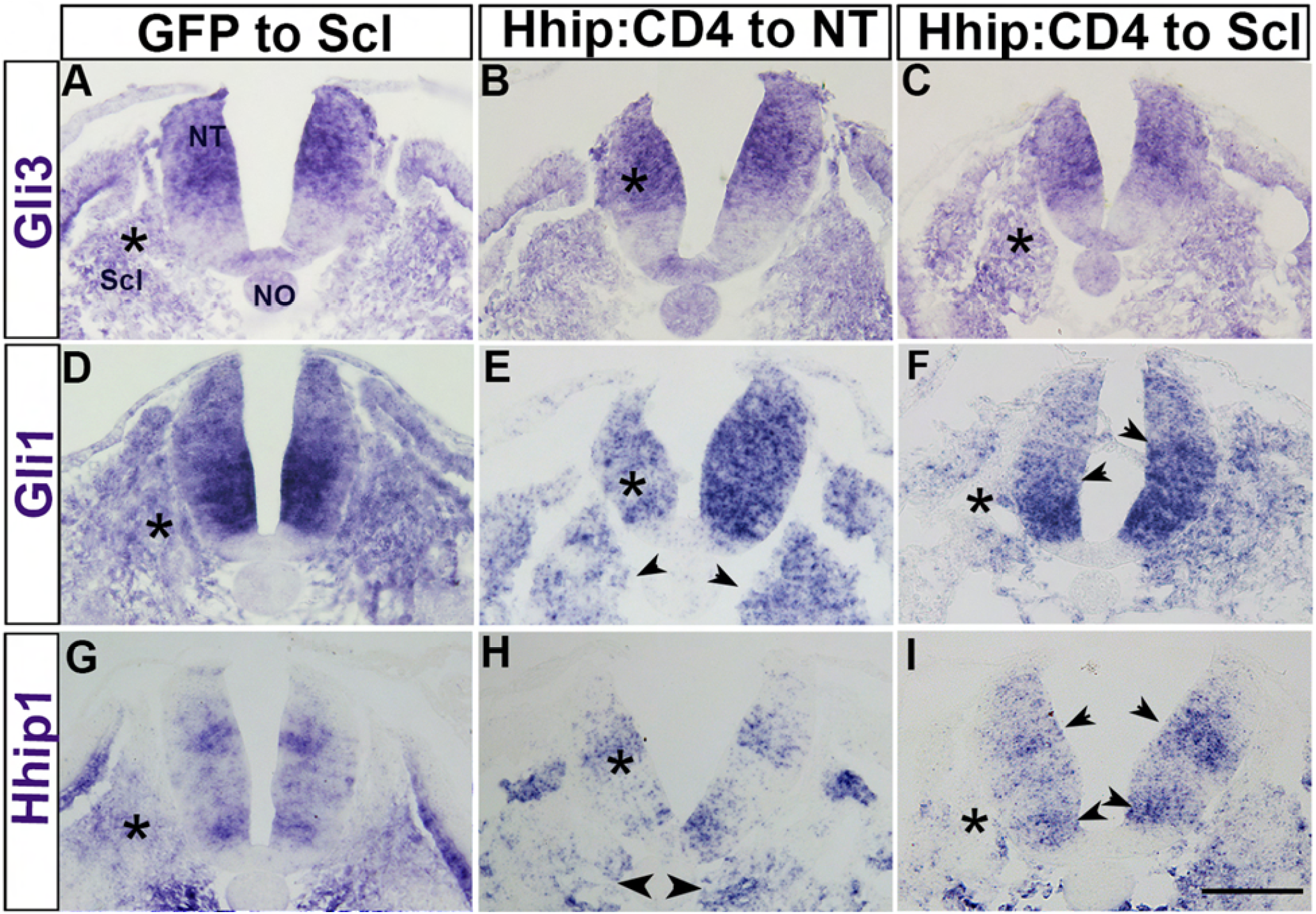
The effects of Hhip:CD4 on NT and myotome are a direct consequence of Shh depletion. (A-I) In situ hybridization for *Gli3, Gli1* and *Hhip1* as depicted following electroporation of control GFP or Hhip:CD4. Asterisks (*) mark the electroporated sites in each panel. Control GFP had no effect on expression of either gene. In contrast, Hhip:CD4 transfected to NT reduced both *Gli1* and *Hhip1* mRNAs in NT and sclerotome (Scl) (arrowheads in E and H). Likewise, Hhip:CD4 transfected to sclerotome reduced both *Gli1* and *Hhip1* mRNAs in NT and sclerotome (Scl) (arrowheads in F and I). No effect was apparent on expression of *Gli3* (A-C). Bar=50 μM.

**Supplementary Fig. S6.**
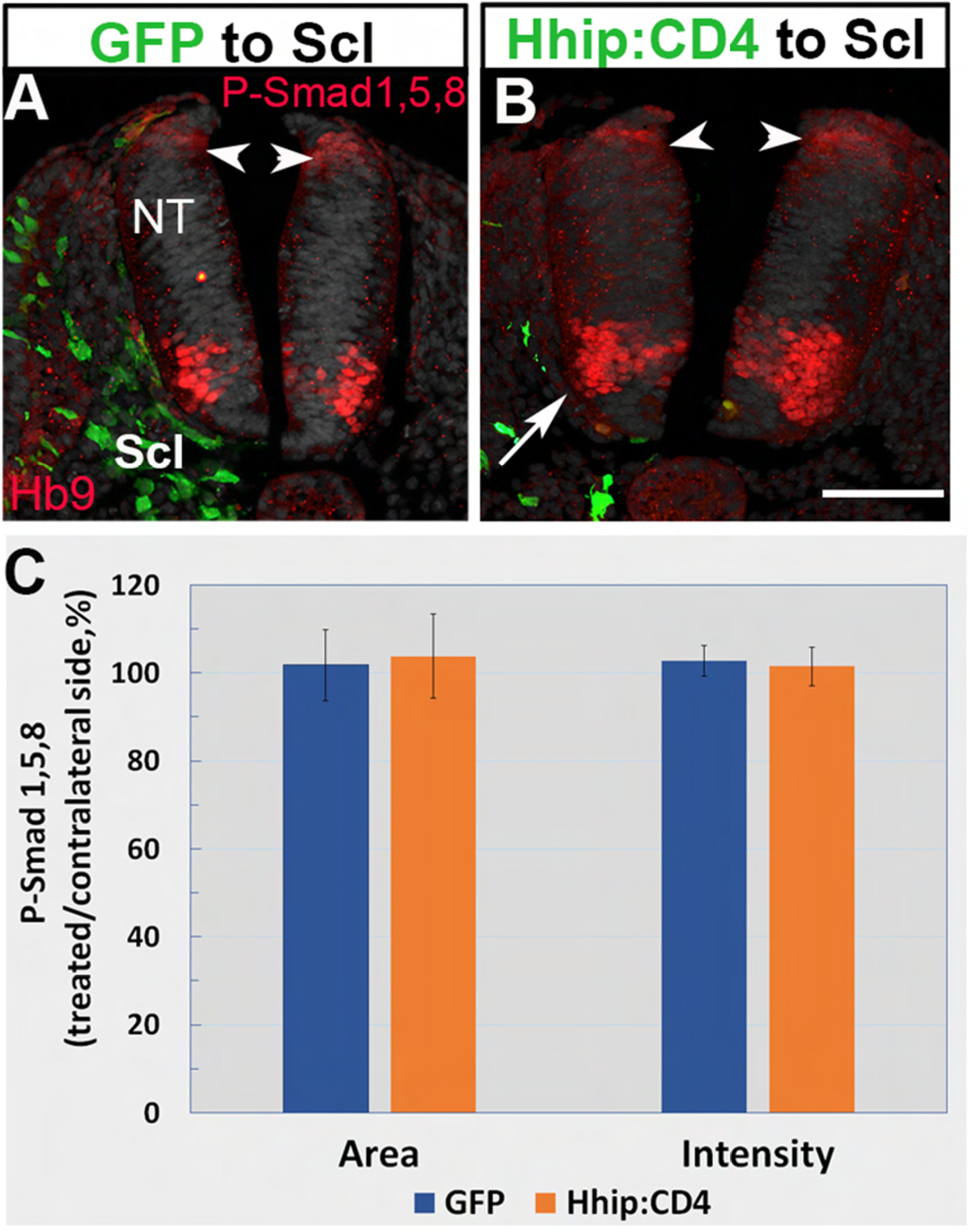
Electroporation of Hhip:CD4 to sclerotome has no effect on BMP signaling in dorsal NT. (A,B) Missexpression of Hhip:CD4 in sclerotome (green) had no effect on expression of phospho-Smad 1,5,8 in the dorsal NT compared to control GFP (arrowheads), yet reduced the number of Hb9+ motoneurons (arrow in B). (C) Quantification of the area and intensity of phospho-Smad 1,5,8 expression. Bar=50 μM.

**Supplementary Fig. S7.**
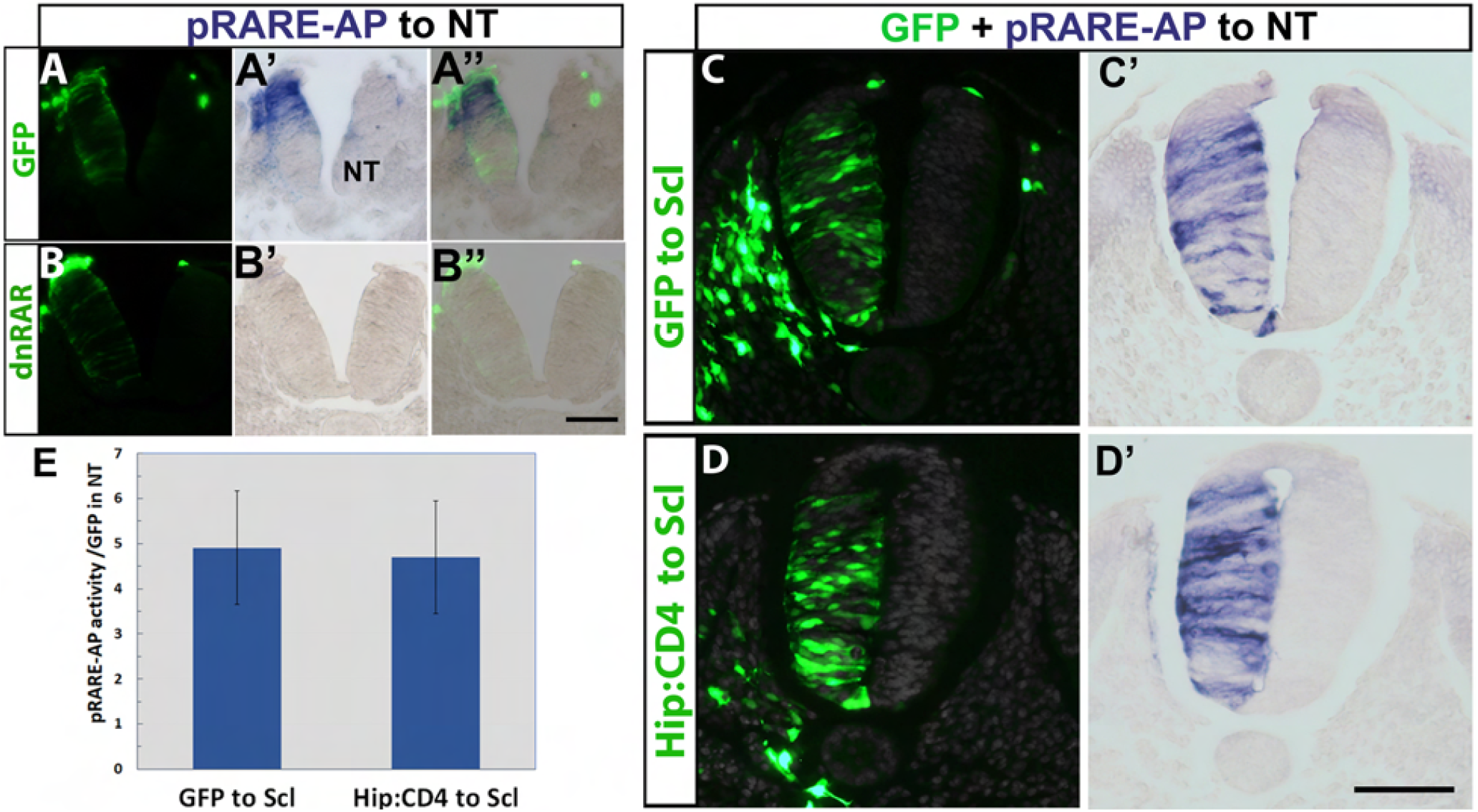
Inhibition of Shh activity in sclerotome has no effect on retinoic acid signaling in the neural tube. (A-B”) Specificity of the retinoic acid reporter (pRARE-AP). Note in A-A” the expression of pRARE-AP in control GFP-electroporated NT. In contrast, no signal is apparent when retinoic acid activity is abolished by a dnRAR plasmid (B-B”). A” and B” are overlays of the precedent panels, respectively. (C-D’) Double electroporations of control GFP or Hhip:CD4 to the sclerotome (C and D, respectively) and co-electroporation of RARE-AP together with GFP to the NT of the same embryos (C,C’, D,D’). (E) Data quantification. Missexpression of Hhip:CD4 in sclerotome had no effect on RARE-AP/GFP activity in NT. Results represent mean values of RARE-AP/GFP±SEM. Bars=50 μM.

